# The p-rpS6-zone delineates wounding response and the healing process

**DOI:** 10.1101/2022.09.27.507541

**Authors:** Nadja Anneliese Ruth Ring, Helene Dworak, Barbara Bachmann, Barbara Schädl, Karla Valdivieso, Tomaz Rozmaric, Patrick Heimel, Kornelia Schuetzenberger, Gabriele Leinfellner, James Ferguson, Susanne Drechsler, Michael Mildner, Markus Schosserer, Paul Slezak, Oded Meyuhas, Florian Gruber, Johannes Grillari, Heinz Redl, Mikolaj Ogrodnik

**Author notes:** These authors contributed equally to this work (Nadja Anneliese Ruth Ring, Helene Dworak).

## Abstract

It is unknown what the spatial boundaries of tissue response to wounding are. Here we show that in mammals the ribosomal protein S6 (rpS6) is phosphorylated in response to skin injury forming a zone of activation surrounding the region of the initial insult. This p-rpS6-zone forms within minutes after wounding and is present until healing is complete. The zone encapsulates markers of the healing process, including proliferation, senescence, and angiogenesis in wounded skin. A mouse model unable to phosphorylate rpS6 shows an initial acceleration of wound closure, but results in disrupted healing. Finally, the p-rpS6-zone accurately reports on the status of dermal vasculature and the effectiveness of healing. In summary, the zone divides an otherwise homogenous tissue into regions with distinct properties.

## Introduction

Wound healing can be divided into three stages: i) the initial damage and its propagation, ii) the immediate molecular response of the injured tissue and iii) healing (Rodrigues et al., 2019). The act of wounding causes damage of tissue and cells resulting in necrosis and apoptosis. This is often followed by an accumulation of debris and bleeding, and leads to rapid changes in ionic composition, oxygen concentration and redox status, breaching of the barrier layers and intrusion of microorganisms to the wounded area (Rodrigues *et al*., 2019). The subsequent early responses of the organism include clotting, vasoconstriction, and mobilization of both immune and non-specialized cells to remove bacteria and debris (Rodrigues *et al*., 2019). Soon after, the regeneration of the damaged or missing tissue structures begins, involving synthesis of extracellular matrix, induction of cellular senescence (Demaria et al., 2014), re-epithelialization and angiogenesis (Rodrigues *et al*., 2019). Owing to technical limitations, a great majority of published research on the immediate reaction to wounding was conducted in indirect models of healing using in vitro 2D cell culture, plants (Toyota et al., 2018), invertebrates (Moreira et al., 2010) and fish (De Simone et al., 2021), with only limited studies performed in mammals (Celli et al., 2021).

Here, we demonstrate that it is possible to visualize and measure the tissue response to wounding as early as minutes and as long as weeks after skin damage, until complete wound closure, using a simple assay to detect a zone of a stable modification of the ribosomal protein S6 (rpS6). RpS6 is an integral component of the 40S subunit of the ribosome and has several phosphorylation sites (S235/236, S240/244 and S247) which are regulated by a range of kinases, including the p70/p85 S6 kinase 1, the p90 ribosomal S6 kinases and protein kinases A, C and G (Biever et al., 2015). The phosphorylation of rpS6 is associated with protein synthesis, cell growth and glucose homeostasis and is increased in the presence of amino acids, glucose and growth factors (Meyuhas, 2015).

The p-rpS6 zone is a new marker of wounding, activated by damage associated molecular patterns (DAMPs) and its induction depends on mTOR signaling pathways, as well as oxygen availability. The p-rpS6-zone encompasses a range of cells involved in skin response to damage, and is characterized by increased proliferation, c-Fos expression, induction of cellular senescence and angiogenesis. Finally, mice that cannot phosphorylate rpS6 show an initial acceleration of healing associated with decreased senescence, but also display a disrupted tissue composition at later stages of healing. Our findings reveal the p-rpS6-zone as an early and stable marker to visualize skin response to wounding. The p-rpS6-zone is a promising prognostic and diagnostic tool to identify the healing status of wounds.

## Results

### Skin damage is delineated by a p-rpS6-zone

Wounding leads to cell and tissue damage, and eventual healing, but the molecular connections between wounding and healing are not well understood. The combination of damaging stimuli and factors causing cell expansion (growth, increase in metabolism and proliferation) can lead to heterogenous cellular responses such as cell death or senescence (Ogrodnik et al., 2019). This concept inspired us to assess changes in the quantity of p-rpS6 level, which is often associated with an increase in the metabolic activity of cells (Meyuhas, 2015) upon wounding and during healing.

We first investigated several injury models in pigs as porcine skin is highly similar to human skin (Sullivan et al., 2001). Specifically, in a burn injury model the skin of animals was wounded using an aluminum cylinder pre-heated to 60 °C, excision injury was performed by taking a 6 mm punch biopsy (scheme in Fig. S1 A) and finally a prick injury was induced using a fine 29-gauge needle (Fig. 1 A). In unwounded control skin the phosphorylation of rpS6 at the sites S235/236 was present solely in the stratum granulosum of epidermis and in some hair follicles (Fig. 1 B). In contrast, at 1.5 h after injury there was a dramatic increase in the amount of p-rpS6 affecting all layers of the skin and cells including keratinocytes, fibroblasts and endothelial cells (Fig. 1 B to H). The signal was visible as an approximately 2 mm thick band matching the orientation of the damage: burn injury led to a horizontal activation zone below the surface, excision injury led to a vertical zone parallel to the damage, while needle injury led to a cylindrical one (consequently smaller in size than for the excision or burn injury) (Fig. 1 B to H). To visualize the dimensionality of the zone in the burn injury, we collected a large skin sample from the edge of the burned area and the subsequent analysis revealed a zone positive for p-rpS6 surrounding the site of the thermal damage (Fig. 1 E and fig. S1 B). In order to visualize the zone around the needle prick sequential tissue slices were cut starting at the epidermis, then stained and reassembled to display the lateral view and the 3D projection of the zone (Fig. 1 H and Movie S1). Finally, we confirmed that the increase in phosphorylation of rpS6 is not limited to serines 235/236, but is also visible when staining for phosphorylation at serines 240/244 using an alternative antibody (Fig. S1 C). Moreover, we were able to validate the increased phosphorylation of rpS6 at serines 235/236 in protein lysate from control, burned and excision-wounded skin by western blot (Fig. S1 D). This suggests that formation of the p-rpS6 zone is a common feature of at least several different types of skin injury: burn, excision and needle injury.

**Fig. 1.**
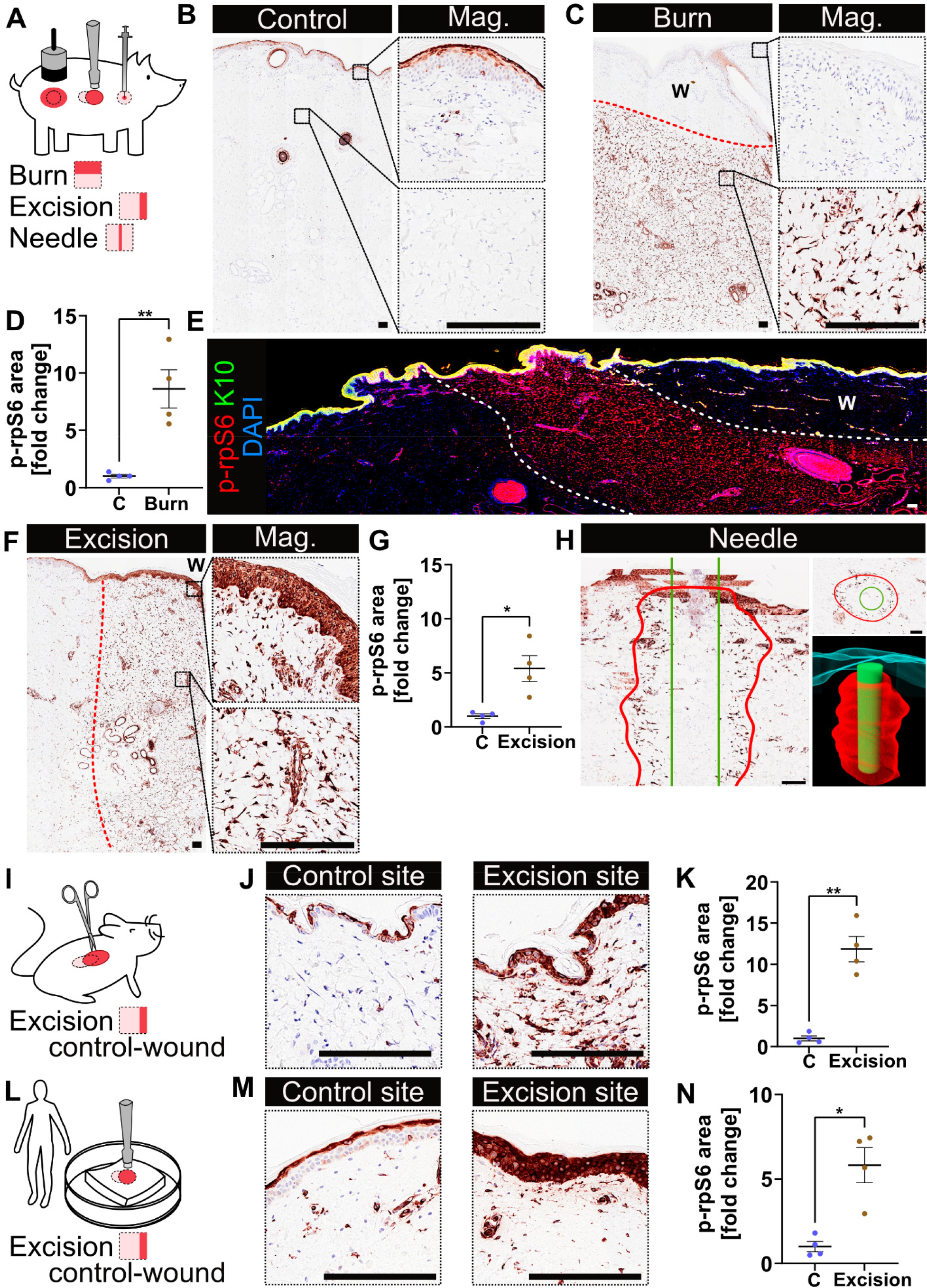
RpS6 phosphorylation marks a zone of tissue activation in response to wounding. Experimental model. Pigs were injured by 60 °C burn, excision wound or needle prick. Samples were collected 1.5 h after injury. (**B**) Porcine skin stained for p-rpS6 from control regions, and (**C**) burn injury, with the edge of the zone marked in red. Micrographs show regions of epidermis and dermis. (**D**) Quantification of p-rpS6 positive area. (**E**) The edge of a 60 °C burn injury, stained with p-rpS6 (red), K10 (green) and DAPI (blue). The burn wound is marked on the top right with a “W”, and the edges of the zone are marked with white dashed lines. (**F**) Porcine excision wound sample stained for p-rpS6, with the edge of the zone marked in red, and the wounded side marked with a “W”. Micrographs show regions of epidermis and dermis. (**G**) Quantification of p-rpS6 positive area. (**H**) Porcine needle prick sample stained for p-rpS6, shown from above (top right panel), in a lateral view by assembling 40 stacked images (left panel), and as a 3D projection (bottom right panel) representing the needle (green) and tissue positive for p-rpS6 staining (red). (**I**) Experimental model in mice. Excision wounds were placed and mice were sacrificed 0.5 - 1.5 h later. (**J**) Murine skin stained for p-rpS6, in regions distal (control site) and adjacent (excision site) to the wound. (**K**) Quantification of p-rpS6 positive area. (**L**) Experimental model in human ex vivo skin. An excision wound was placed using a biopsy punch, and samples were collected from the edge of the wound after 1.5 h. (**M**) Human *ex vivo* skin stained for p-rpS6 in regions proximal (excision site) and distal (control site) to the wound site. (**N**) Quantification of p-rpS6 positive area. Data are from n = 4 pigs per group for (D) and (G); n = 4 mice per group for (K); n = 4 human subjects for (n). Mean ± SEM plotted. For all graphs unpaired t-test was used. *p<0.05, **p<0.01. The scale bars for all the images are 100 μm.

### The p-rpS6-zone is conserved in mammals

To assess how reliable and evolutionarily-conserved the induction of the p-rpS6-zone is, we used murine skin injured by full-thickness excision wound. Mice were sacrificed 0.5 to 1.5 h after injury (Fig. 1 I). Matching our observations in porcine skin, we found a thin layer of p-rpS6 limited to the stratum granulosum of unwounded skin, while wounded murine skin displayed a broad zone of activated rpS6 (Fig. 1 J). Similar to porcine skin, the p-rpS6-zone in murine skin was approximately 1.5 mm wide and encompassed a variety of cell types in all skin layers (Fig. 1 J, K and Fig. S1 E). Finally, to determine the value of these findings for humans, we used human skin collected as a by-product of surgical procedures (Fig. 1 L). We performed excision wounds using a 6 mm punch biopsy in the skin within an ex vivo setting and again found a strong induction of the p-rpS6-zone (Fig. 1 M, N and fig. S1 F), confirming evolutionary conservation of this response across mammals (Fig. 1 A to N).

### The zone stratifies burns

Clinical practice involves the assessment of wound severity using measurements of cell death, predominantly necrosis and apoptosis (Holzer et al., 2020). However, to our knowledge there are currently no reliable tissue markers that could be used to describe the range of tissue response to wounding without relating directly to cell death. Intrigued by our earlier observation that the p-rpS6-zone is directly adjacent to the damage in an excision wound and instead appears 2 mm below the skin surface in burn wounds (Fig. 1 C and F), we hypothesized about the relationship between p-rpS6 induction and cell death. Thus, we investigated if cell death by heat exposure might be responsible for the distinctive location of the zone in burn wounds.

We used two well established markers of cell death: high mobility group box 1 (HMGB1) – a protein which leaks out of the nucleus to the cytoplasm in response to cell stress and necrosis, and cleaved caspase 3 – a marker of apoptosis. We found that at 1.5 h after a 60 °C burn, the cells in the top 2 mm of the skin displayed HMGB1 leakage while the HMGB1 content of the cells below that layer remained unaffected (Fig. 2 A and D). Cells positive for cleaved caspase 3 were present in a thin layer of approximately 0.5 mm width that overlapped with the lower part of the necrotic layer (Fig. 2 B and D).

**Fig. 2.**
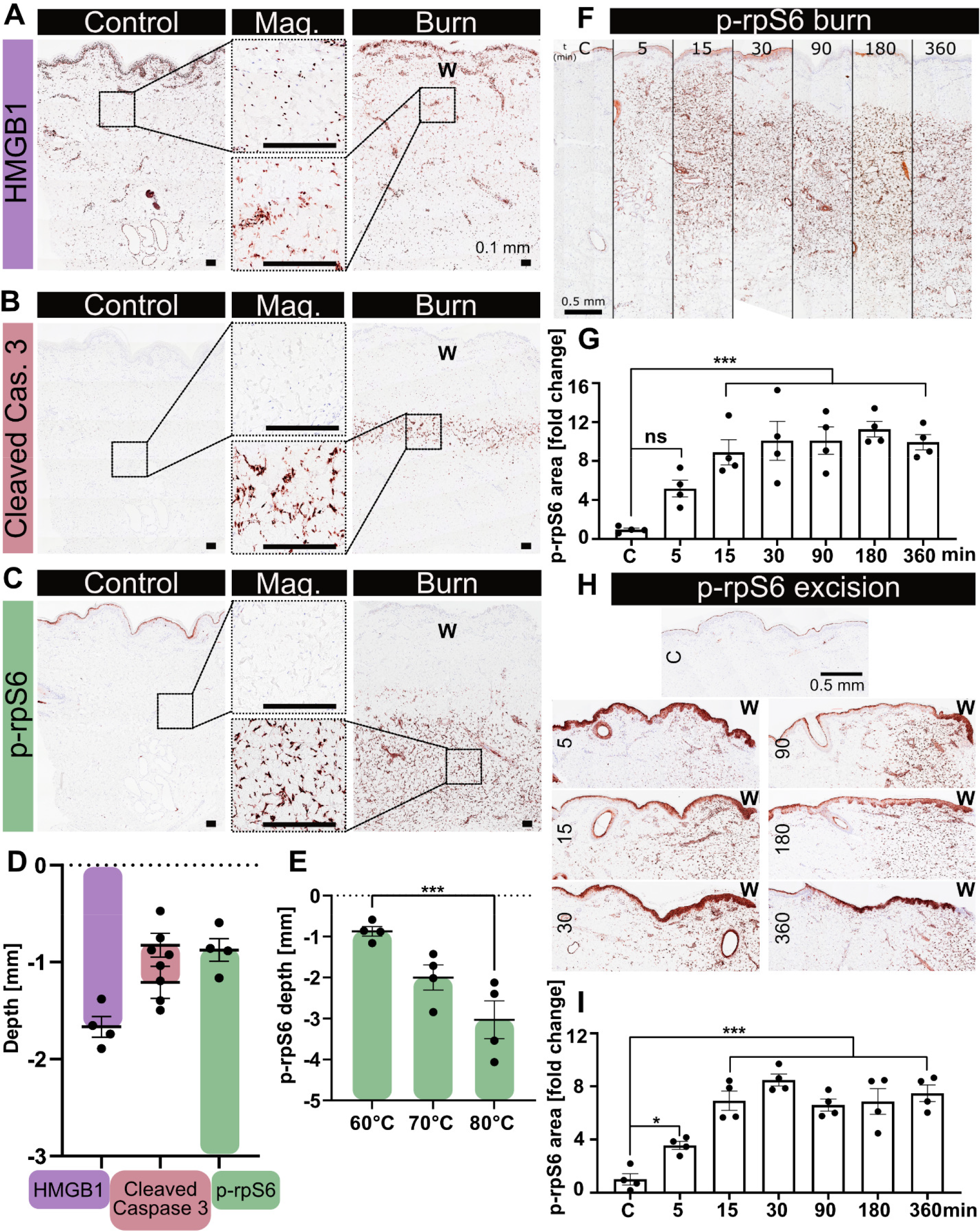
The induction of the p-rpS6-zone is an early-onset and long-term response that stratifies cell death and survival responses. (**A**) Porcine skin stained for HMGB1, (**B**) cleaved caspase 3, and (**C**) p-rpS6 in control and 60 °C burn injury (burn wound is marked with a “W”). Micrographs show magnified regions of dermis. (**D**) Quantification of the depth and thickness of the responses to a 60 °C burn injury at 1.5 h post-injury. Purple marks the thickness of layer of HMGB1 leakage, red shows the depth of initiation (top datapoints) and thickness (bottom datapoints) of the cleaved caspase 3 layer, while green shows the depth of initiation of the p-rpS6 layer. (**E**) Quantification of the depth of initiation of the p-rpS6-zone at 1.5 h after a 60, 70 and 80 °C burn injury. (**F**) Porcine skin stained for p-rpS6 in control conditions and several timepoints after 60 °C burn. (**G**) Quantification of p-rpS6 positive area. (**H**) Porcine skin stained for p-rpS6 in control conditions several time points after excision wound (the wound is on the right side of the biopsy and marked with a “W”). (**I**) Quantification of p-rpS6 positive area. Data are from n = 4 pigs per group. Mean ± SEM plotted. For (E), (G) and (I) one-way ANOVA with post-hoc Dunnet’s test was used. The scale bars for (A), (B) and (C) show 100 μm; for (F) and (H) show 500 μm. *p<0.05, ***p<0.001 and “ns” is “non-significant”.

The p-rpS6-zone starts with the beginning of the apoptotic layer, and stretches deep into the dermis, beyond all markers of cell death (Fig. 2 C and D). With higher temperatures of 70 and 80 °C, we observed a downward shift of both the HMGB1 layer (Fig. S2 A, B) and the p-rpS6-zone (Fig. 2 E and fig. S2 C) by ∼1 mm per 10 °C. This suggests that this stratification relates directly to burn-induced skin response, with higher temperatures inducing an increased extent of cell death and therefore an initiation of the p-rpS6-zone which is deeper in the skin. The p-rpS6 signal present in the stratum corneum of control tissue is obliterated by the burn, probably due to burn- and necrosis-induced protein degradation or dephosphorylation. In contrast, 1.5 h after excision wounding, we observed no evident leakage of HMGB1 adjacent to the wound suggesting an insignificant amount of cell death, while the p-rpS6-zone was strongly induced (Fig. S2 D and E). These results demonstrate that the p-rpS6-zone is a novel and powerful marker of tissue response to wounding, which appears in response to damage without being specific to dying cells. The p-rpS6-zone potentially constitutes the first stable early-onset damage-indued marker that goes beyond cell death.

### The zone is immediate and long-lasting

With the majority of markers related to tissue wounding – such as infiltration of immune cells (Rodrigues *et al*., 2019) – appearing hours to days after induction of damage, we sought to determine how rapidly and stably p-rpS6 is induced in excision and burn injuries. One exception is the Erk signaling pathway, which was recently found to be induced in a spatially-controlled manner in response to wounds in invertebrates and fish (De Simone *et al*., 2021). In order to compare the time of induction and the propagation dynamics of p-rpS6 and p-Erk we analyzed samples at minutes to hours after burn or excision injury in a porcine model (Fig. 2).

Burn injury resulted in the appearance of the p-rpS6-zone within minutes and reached statistical significance at 15 min after wounding (Fig. 2 F, G). Moreover, the intensity of the signal plateaued and remained present in high quantity and of significant intensity even at 6 h post-injury (Fig. 2 F, G). When analyzing the depth of initiation and termination of the p-rpS6-zone, we did not observe any significant changes in the depth of initiation within 6 h after injury (Fig. 2 F and fig. S3 A, B). However, we did observe a gradual propagation of the terminus of the p-rpS6-zone, which penetrated deeper into dermis over time, an effect especially visible at 6 h (Fig. 2 F and fig. S3 A, C).

Similar to p-rpS6, p-Erk was detectable very early after burn injury, and indeed p-Erk had faster dynamics than p-rpS6, showing a sharp and significant rise as early as 5 min after injury (Fig. S3 D and E). However, the rapid increase was followed by an equally rapid decrease and 30 min after injury the level of p-Erk was no longer significantly different from control (Fig. S3 E) indicating an acute induction of p-Erk with a rapid decline. Unlike p-rpS6, which showed a spatially discrete stratification (Fig. 1 and fig. 2), the p-Erk signal was diffuse, with no obvious changes in propagation over time (Fig. S3 D).

Consistently, excision wounds also showed an early and long-lasting induction of p-rpS6 reaching significance at 5 min after wounding and remaining elevated throughout the entire monitoring period (Fig. 2 H, I). In contrast to burn injury, however, we did not detect any significant changes in the spatial positioning of the p-rpS6-zone, which stably marked a layer of around 2 mm from the side of the excision wound (Fig. 2 H and fig. S3 F, G).

Just as in burns, the significant increase of p-Erk signal in excision wounds was detectable as early as 5 min after injury (Fig. S3 H and I). However, in contrast to p-rpS6 the statistical significance was lost for timepoints collected after 30 min of injury and unlike p-rpS6, we did not observe any consistent spatial relationship to the wound site (Fig. S3 H and I).

While HMGB1 leakage occurred as early as 5 min after induction of the burn wound, there was no significant propagation of the necrotic zone within the time period of up to 6 h post injury (Fig. 3 J). This suggests that the p-rpS6-zone appears quickly and concomitantly with the earliest markers of cell death, and is stable, and spatially discrete; the p-rpS6-zone therefore represents a fast and easy to perform technique to detect and quantify damage response at tissue level.

**Fig. 3.**
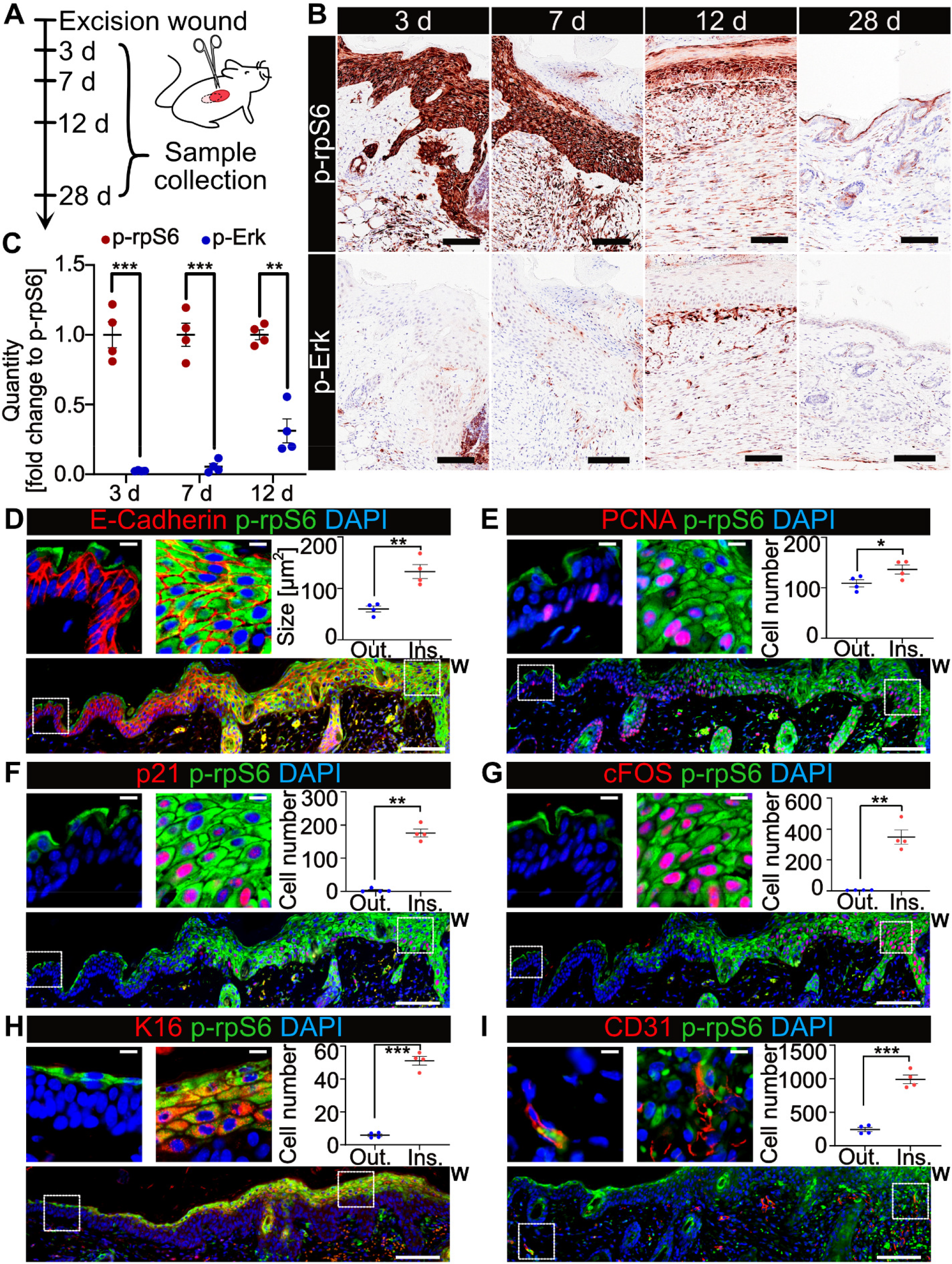
The p-rpS6-zone is present throughout the healing process and encompasses pro-healing cellular processes. (**A**) Experimental model in mice. Excision wounds were placed and mice were sacrificed 3, 7, 12 or 28 days later. (**B**) Murine skin stained for p-rpS6 and p-ERK at several time points after excision wound. (**C**) Quantification of p-rpS6 and p-ERK area. (**D-G**) Murine skin at 3 d post injury, with micrographs and quantifications of regions outside (Out.) or inside (Ins.) the p-rpS6-zone; the zone is immediately adjacent to the wound (marked with a “W”). (D) E-cadherin (red), p-rpS6 (green) and DAPI (blue). Quantification of average size of keratinocytes, 150-200 cells per mouse were quantified. (E) PCNA (red), p-rpS6 (green) and DAPI (blue). Quantification of average number of PCNA positive keratinocytes, 5 images per mouse were quantified. (F) p21 (red), p-rpS6 (green) and DAPI (blue). Quantification of average number of p21-positive keratinocytes, 5 images per mouse were quantified. (G) cFos (red), p-rpS6 (green) and DAPI (blue). Quantification of average number of cFos-positive keratinocytes, 5 images per mouse were quantified. (**H-I**) Murine skin at 12 d post injury. (H) K16 (red), p-rpS6 (green) and DAPI (blue). Quantification of average number of K16-positive keratinocytes, 6 images per mouse were quantified. (I) CD31 (red), p-rpS6 (green) and DAPI (blue). Quantification of average number of CD31-positive cells in the dermis, normalized to the area of the dermis, 14 images per mouse were quantified. Data are from n = 4 mice per group for all the graphs. Mean ± SEM plotted. For the graph (C) multiple unpaired t-test was used (false discovery rate >1 %). For (d-i) paired t test was used. *p<0.05, **p<0.01, ***p<0.001. Scale bars for (B) and large images in (D-I) show 100 μm, for the micrographs in (D-I) scale bars are 10 μm.

Our results univocally demonstrate that the p-rpS6-zone is an early-response and stable marker of tissue injury. As our results on the rapid appearance of the p-rpS6-zone come from in vivo porcine skin wounds as well as ex vivo human skin wounds, these findings are directly translatable to humans.

### The zone encompasses healing processes

The process of wound healing is associated with a variety of histological features including an increased proliferation and size of keratinocytes (Rodrigues *et al*., 2019), induction of cellular senescence (Demaria *et al*., 2014), expression of proto-oncogenes such as c-Fos (Martin and Nobes, 1992), infiltration of immune cells and angiogenesis (Rodrigues *et al*., 2019). However, the common molecular traits connecting these features are not well established.

In order to investigate the relevance of the p-rpS6-zone in the healing process we analyzed murine skin samples at 3, 7, 12 and 28 days after wounding following the scheme shown in Fig. 3 A. In our experiment, wounds were re-epithelialized by day 12 and completely healed by day 28 (Fig. S4 A). The p-rpS6-zone was detectable throughout the process of healing and the p-rpS6 signal was present in keratinocytes forming the epithelial tongue, in dermal fibroblasts proximal to the injury site and in re-growing blood vessels (representative images in Fig. 3 B are showing the wound edge as demonstrated in Fig. S1 E). The zone was lost after completion of the healing process (Fig. 3 B), with p-rpS6 levels returning to control levels in both epidermis and dermis (Fig. 1). We observed that just as in the case of p-rpS6 (Fig. 1), wounding of murine skin induces phosphorylation of Erk within a short period of time in all skin layers (Fig. S4 B). However, p-Erk was nearly absent in wounded or wound-proximal skin at 3, 7 and 12 days after wounding (Fig. 3 B, C), which is consistent with the findings on the declining p-Erk levels within hours from injury in pigs (Fig. 3) and suggests that its induction is transient and diminishes shortly after wounding. Interestingly, the highest level of p-Erk was observed at day 12 in periepidermal fibroblasts (Fig. 3 B and C), which roughly corresponds to the time when fibroblasts are eliminated by immune cells or undergo apoptosis (Rodrigues *et al*., 2019).

Next, we compared features of cells found inside the p-rpS6-zone with those in homeostatic skin immediately proximal to the zone. We found that cells positive for markers canonically related to the process of re-epithelialization and associated with epidermal thickening, such as an increase in keratinocyte size (Fig. 3 D) and increased keratinocyte proliferation (Fig. 3 E) are upregulated within the p-rpS6-zone. In respect to proliferation, we found that ∼95% of keratinocytes positive for PCNA (a marker of proliferation) were also positive for p-rpS6 (Fig. S4 C). Moreover, we found that a commonly used marker of cellular senescence (p21CIP1/WAF1, Cdkn1a) was present nearly exclusively in keratinocytes within the p-rpS6-zone (Fig. 3 F and fig. S4 C). We confirmed the enrichment in markers of senescence within the p-rpS6-zone by staining for irreparable DNA double strand breaks (DSBs) at telomeres, telomere associated foci (TAF), a reliable marker of senescence (Hewitt et al., 2012). This staining is performed by pairing an immunostaining against γ-H2A.X, a marker of DSBs, with a TelC probe to detect telomeric sequences. Both the total number of cells positive for γ-H2A.X and TAF were highly increased in keratinocytes within the p-rpS6-zone when compared to keratinocytes immediately outside of the zone (Fig. S4 D).

Similarly, the proto-oncogene c-Fos, which is involved in the mechanisms of healing-related differentiation and migration of keratinocytes (Martin and Nobes, 1992) was also found to be present almost exclusively in keratinocytes positive for p-rpS6 (Fig. 3 G and fig. S4 C). Another marker of re-epithelialization, expression of keratin 16 (K16) was present at 12 days after wounding and was almost exclusively inside the p-rpS6-zone and p-rpS6 positive (Fig. 3 H and fig. S4 C). In contrast, CD3-positive T cells present in the dermal compartment of the wounded area were found to be predominantly negative for p-rpS6 (Fig. S4 C and E). Another essential marker of healing is angiogenesis, which can be characterized by an increase in density of blood vessels (Rodrigues *et al*., 2019) and appearance of endothelial tip cells, which express proteins such as endocan (also known as endothelial cell-specific molecule 1; Esm1) (Rocha et al., 2014). Consistently, we found that the dermal compartment of the p-rpS6-zone is enriched in blood vessels (Fig. 3 I), and is specifically enriched in endocan-positive tip cells, most of which are also p-rpS6 positive (Fig. S4 C, F).

In summary, our data from murine wounds indicate that the p-rpS6-zone is not only related to the immediate response to wounding, but is also present throughout the whole process of healing while being associated with re-epithelialization, induction of cellular senescence, c-Fos expression and angiogenesis.

### Zone formation can be modelled *in vitro*

The S235/236 phosphorylation site of rpS6, which we primarily focused on to define the p-rpS6-zone, can be phosphorylated by several proteins including S6K1 and 2, RSK1 and PKC (Biever *et al*., 2015) linking it to several master regulators of cell physiology including mTOR and Erk.

To unravel the mechanism leading to the formation of the damage-induced p-rpS6-zone, we used a range of chemical compounds to manipulate key players in a variety of signaling pathways in an in vitro scratch assay using primary human dermal fibroblasts (HDFs). High levels of p-rpS6 are constitutively present in cell culture conditions due to the presence of fetal bovine serum (FBS) in cell culture media (Fig. S5 A). However, a short incubation period with basal media lacking FBS (basal media) reduced the overall level of p-rpS6 while preserving normal morphology of cells (Fig. S5 A). In these conditions, we observed that induction of a scratch results in a significant elevation of p-rpS6 in cells proximal to the damaged area forming a zone, reminiscent of that found in vivo (Fig. 4 A, B and fig. S5 B). Consistent with experiments on live pigs (Fig. 2), induction of p-rpS6 occurs shortly after execution of the scratch and persists for a prolonged period of time (Fig. 4 A, B; and fig. S5 B). We further confirmed the scratch-dependent increase in p-rpS6 in the proximity to the insult in a model of keratinocytes, using HaCat, a spontaneously immortalized human keratinocyte cell line (Fig. S5 C).

**Fig. 4.**
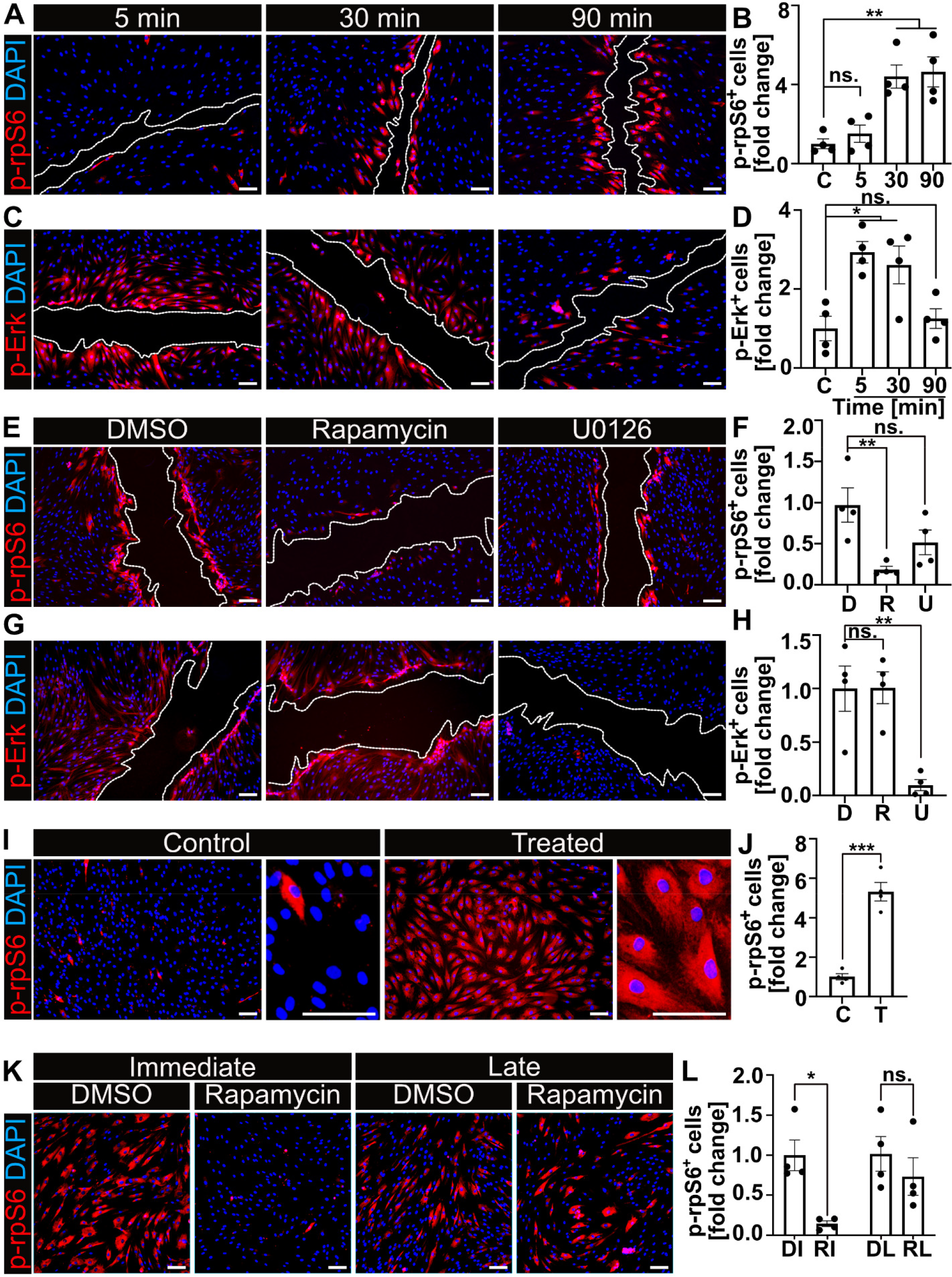
Formation of the p-rpS6-zone is dependent on mTOR and induced by DAMPs. (**A**) Human dermal fibroblasts (HDFs) fixed 5, 30 or 90 min after scratch and stained for p-rpS6 (red) and DAPI (blue). (**B**) Quantification of the number of p-rpS6-positive cells (fold over unscratched control cells, “C”). (**C**) HDFs at 5, 30, or 90 min post-scratch, stained for p-Erk (red) and DAPI (blue). (**D**) Quantification of the number of p-ERK-positive cells (fold over control). (**E**) HDFs treated with DMSO (“D), rapamycin (“R”) or U0126 (“U”) and fixed 30 min post-scratch, stained for p-rpS6 (red) and DAPI (blue). (**F**) Quantification of the number of p-rpS6-positive cells (fold over DMSO control). (**G**) HDFs treated as in panel e and stained for p-Erk (red) and DAPI (blue). (**H**) Quantification as in panel f. (**I**) HDFs left untreated (control, “C”) or treated with media collected from mechanically lacerated cells (“T”) and fixed after 30 min, stained for p-rpS6 (red) and DAPI (blue). (**J**) Quantification of the number of p-rpS6-positive cells (fold over control). (**K**) HDFs treated as in panel i, and co-treated with vehicle (DMSO, “D”) or rapamycin (“R”) either immediately (“I”) or 30 min later (“L”). All cells were fixed 30 min after administration of the drug. (**L**) Quantification of the number of p-rpS6-positive cells (fold over control, immediate DMSO treatment “DI”). Data are from n = 4 biological replicates (independent experiments) per group for all the graphs. For all quantifications 10 images per biological replica were used. For all the graphs mean ± SEM plotted. For the graphs (B), (D), (F), (H) and (J) one-way ANOVA with post-hoc Dunnet’s test was used. For (L) two-way ANOVA with post-hoc Sidak’s test was used. *p<0.05, **p<0.01, ***p<0.001. All scale bars show 100 μm.

In HDFs the scratch assay also induced an increase in p-Erk: while it occurred earlier than rpS6 activation, at 5 min after scratching, its levels declined over time and returned to the control level within 90 min (Fig. 4 C, D and fig. S5 D). The assay thus sufficiently mirrors the early response to wounding in vivo and can be used to study pathways leading to the phosphorylation of rpS6.

### Zone formation is mTOR dependent

Pre-treatment of cells with rapamycin, a potent inhibitor of mTOR, resulted in an almost complete abrogation of the p-rpS6 induction in the scratch assay, but did not affect Erk phosphorylation. In contrast, the use of U0126 (an inhibitor of MEK, the upstream regulator of Erk) had no significant effect on rpS6 phosphorylation (Fig. 4 E, F and fig. S5 E) while almost completely abrogating p-Erk-positive cells (Fig. 4 G, H and fig. S5 F). These results therefore suggest that the induction p-rpS6 and p-Erk are two independent events regulated separately by the upstream master regulators mTOR and MEK respectively.

### Zone formation is induced by DAMPs

Factors released immediately after wounding are grouped under the term damage-associated molecular patterns (DAMPs), and include proteins, lipids, nucleic acids and small compounds such as ATP or glutamate. While these normally reside inside cells and their organelles, they are released or passively leaked from damaged and dying cells (Cordeiro and Jacinto, 2013). In order to establish whether DAMPs induce formation of the p-rpS6-zone in response to wounding, or if it is the wounded cells themselves which respond, we performed a “passive scratch assay”. Specifically, donor cells were lacerated using a needle (i.e. scratch assay was performed in the whole area of the culture vessel), releasing DAMPs into the cell culture media. This DAMP-containing media was then directly transferred to non-scratched recipient cells, which were fixed after 30 min and stained. We found that the presence of DAMPs in culture media causes a striking and homogenous population of p-rpS6-positive recipient cells throughout the cell culture plate (Fig. 4 I and J).

To assess whether a functional mTOR complex is needed for the induction or maintenance of the p-rpS6-zone we either treated cells with rapamycin immediately prior to the addition of DAMPs or treated them with rapamycin later when the p-rpS6-zone was fully established. For both modes of treatment cells were incubated with the drug for 30 min. Surprisingly, we found that when rapamycin is added at the beginning of the assay, it prevents formation of the p-rpS6-zone when compared to the control (DI and RI; DMSO immediate and rapamycin immediate, respectively) while the same duration of time with the drug added after zone formation did not significantly affect the frequency of the p-rpS6-positive cells (DL and RL; DMSO late and rapamycin late, respectively) (Fig. 4 K and L). Overall, these results suggest that DAMPs are causal to mTOR-mediated phosphorylation of rpS6 in cells proximal to the damage. However, it remains unclear at this point how the phosphorylation status of rpS6 is affecting the progression of healing.

### P-rpS6 deficiency accelerates wound closure

To assess the role of the p-rpS6-zone in healing, we used a mouse model which is constitutively unable to phosphorylate rpS6. In these rpS6 p^−/-^ KI (rpS6 KI) mice, the serine residues 235, 236, 240, 244, and 247 of rpS6 have been replaced with alanines through site-directed mutagenesis, making them unable to phosphorylate this protein (Ruvinsky et al., 2005). Comparing rpS6 KI to wild-type C57bl/6 (WT), we performed a wounding assay (Fig. S6 A) consistent with our earlier experiments (Fig. 3), with mice of both genotypes body weight matched (Fig. S6 B). We discovered that the wound closure of rpS6 KI mice is significantly accelerated at the very early stages of wound healing in females (Fig. 5 A, B and fig. S6 A) and to a lesser degree in males (Fig. S6 C). Upon sacrifice, we verified the phospho-deficient genotype of all individual mice by IHC staining, by confirming the lack of p-rpS6 in the KI mice (representative image shown in fig. S6 D). The skin tissue from mice sacrificed at day 3 post-wounding revealed a decline in cellular senescence as denoted by a significant decrease in cells positive for p21 in rpS6 KI mice (Fig. 5 C and D). Correspondingly, the number of cells positive for the proliferation marker PCNA showed a slight increase in the rpS6 KI mice (Fig. 5 E and F). In accordance with published results from in vitro experiments (Ruvinsky *et al*., 2005), we found that the wounding-induced increase in the size of keratinocytes is diminished in skin of in the rpS6 KI mice (Fig. 5 G and H). In contrast, we did not observe differences in the number of c-Fos positive cells (Fig. S6 E and F). These results indicate that the p-rpS6-zone may be driving senescence, inhibiting proliferation and increasing cell size in the tissue surrounding a wound. While the inhibition of this phosphorylation may speed up the initial phases of wound healing, we were curious to investigate the long-term consequences of the lack of the p-rpS6-zone.

**Fig. 5.**
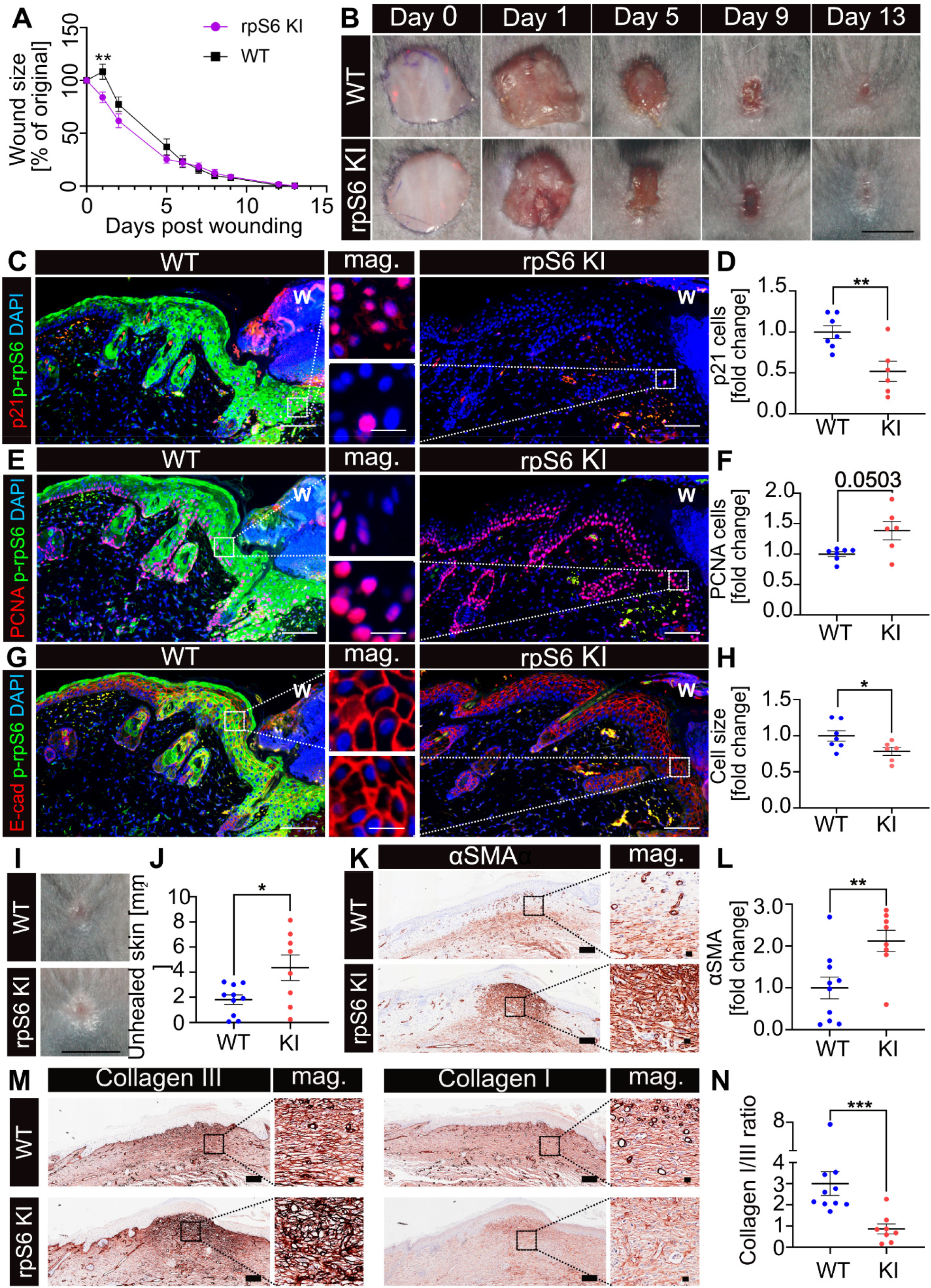
P-rpS6 deficiency results in faster initial wound closure but disrupted healing. rpS6 KI and WT mouse excision wounds were photographed at days 0, 1, 2, 5, 6, 7, 8, 9, 12 and 13. Wound size was normalized to the measurement from day 0 and data is shown as percentage of initial size. (**B**) Representative photographs of wounds from rpS6 KI and WT mice. (**C-H**) Murine skin at 3 d post injury (wounded side marked with a “W”), p-rpS6 staining has been excluded from all micrographs for clarity. Quantifications of the regions immediately adjacent to the wound in rpS6 KI and WT mice. (C) p21 (red), p-rpS6 (green) and DAPI (blue). (D) Quantification of average number of p21-positive keratinocytes. (E) PCNA (red), p-rpS6 (green) and DAPI (blue). (F) Quantification of average number of PCNA positive keratinocytes. (G) E-cadherin (red), p-rpS6 (green) and DAPI (blue). (H) Quantification of average size of keratinocytes, 100-200 cells per mouse were quantified. (**I**) Representative images comparing healing progression at day 13, in WT and rpS6 KI mice. (**J**) Quantification of the macroscopically-visible region of light hairless skin. (**K**) Murine skin from WT and rpS6 mice at 13 d post injury and stained for αSMA. (**L**) Quantification of αSMA positive area. (**M**) Murine skin from WT and rpS6 mice at 13 d post injury and stained for collagen III (left side) and collagen I (right side). (**N**) The ratio of collagen I to collagen III in skin samples of WT and rpS6 KI mice. Data are from n = 8 rpS6 KI and n = 10 WT female mice for the graphs (A), (J), (L) and (N) and from n = 6 rpS6 and n = 7 WT male mice for the graphs (D), (F) and (H). Mean ± SEM plotted for all the graphs. For the graph (A) two-way ANOVA with post-hoc Sidak’s test was used. For (D), (F), (H), (J) and (L) unpaired t test was used. For (N) Mann-Whitney U test was used. *p<0.05, **p<0.01 and ***p<0.001. Scale bars for the photographs in panels (B) and (I) are 5 mm. Scale bar for the images at (C), (E), (G), (K) and (M) show 100 μm and for the micrographs 10 μm.

### P-rpS6 deficiency results in disrupted healing

While the wound closure of the rpS6 KI mice matched the kinetics of WT mice, with near complete closure by day 13 post-injury, there were clear differences visible both macroscopically and histologically (Fig. 5 B and I). Macroscopically, 13 days after injury many of the wounds of rpS6 KI mice are outlined by large rings of light-colored and hairless skin, indicating an incomplete and disrupted healing process (Fig. 5 B, I, J and fig. S6 A). Histologically, the final steps of wound healing involve reduction of alpha smooth muscle actin (αSMA) and replacement of collagen III by collagen I (Rodrigues *et al*., 2019). In contrast, histological analysis revealed that late-stage wounds of rpS6 KI mice show a significant accumulation of αSMA (Fig. 5 K and L). In addition, the wounds of rpS6 KI mice have comparatively more collagen III, less collagen I, and a lower collagen I/III ratio (Fig. 5 M, N and fig. S6 G, H). Altogether, these results demonstrate that interfering with the p-rpS6-zone can lead to an acceleration of healing at an early stage, but results in disrupted healing at later stages, suggesting that the p-rpS6-zone is important for the timely resolution of a wound. We next sought to explore pathophysiological scenarios where the p-rpS6-zone may be similarly dysregulated.

### Hypoxia prevents p-rpS6-zone formation

Lack of circulation (ischemia) and the resulting lack of oxygen (hypoxia) is at the heart of many types of chronic wounds including venous, pressure and diabetic ulcers (Cox, 2013; Han and Ceilley, 2017; Zhao et al., 2016). It was also shown that hypoxia reduces the activity of mTOR (Brugarolas et al., 2004; van Vliet et al., 2021). We therefore sought to determine whether the hypoxic conditions resulting from a loss of circulation could affect the formation of the p-rpS6-zone resulting from skin injury, mimicking the lack of p-rpS6 seen in the phospho-deficient mouse model used above.

To test this hypothesis, we performed burn and excision injuries on alive and sacrificed pigs – as burn injuries do not breach the skin surface they remain hypoxic when circulation is halted after death, while the tissue of excision injuries is exposed to the surrounding air and is never fully hypoxic. Three types of porcine skin biopsy samples were collected: (i) a set of injuries was performed on anesthetized animals and the samples were collected after 30 min (“alive”), (ii) a second set of injuries was performed simultaneously, the animals were sacrificed after 30 min and the samples were collected 30 min post-sacrifice (60 min post injury, “alive-dead”), (iii) a final set of injuries was performed shortly after sacrifice and samples were collected after 30 min (“dead”; Fig. 6 A). The lack of circulation in the dead animal was macroscopically visible as the burn induced no erythema and the excision wound caused no bleeding (Fig. S7 A), while skin surface temperature, which was measured prior, during and after burn changed only by 1-2 degrees (Fig. S7 B). Importantly, we noted large differences in the level of p-rpS6 induction (Fig. 6 B and C). In fact, while burn injuries from live animals showed a large quantity of p-rpS6 below the necrotic zone, the induction of the p-rpS6-zone was completely abrogated in burn samples collected from the recently-deceased animals (Fig. 6 B and C). Surprisingly, the “alive-dead” samples showed no significant difference in the levels of p-rpS6 when compared to samples collected from alive animals (Fig. 6 B and C). In contrast to the sharp decline in p-rpS6 levels in the burn wounds of recently-deceased animals, the excision wounds did not show any significant differences in p-rpS6 levels between samples collected from alive, dead and “alive-dead” conditions (Fig. 6 D and E). We hypothesize that the difference in the response of the skin of the dead animals to the excision and burn wounds is due to the fact that an excision wound is exposed to atmospheric oxygen, allowing the regions nearest to the wound to activate the p-rpS6-zone, even when oxygen from the blood vessels is no longer circulating.

**Fig. 6.**
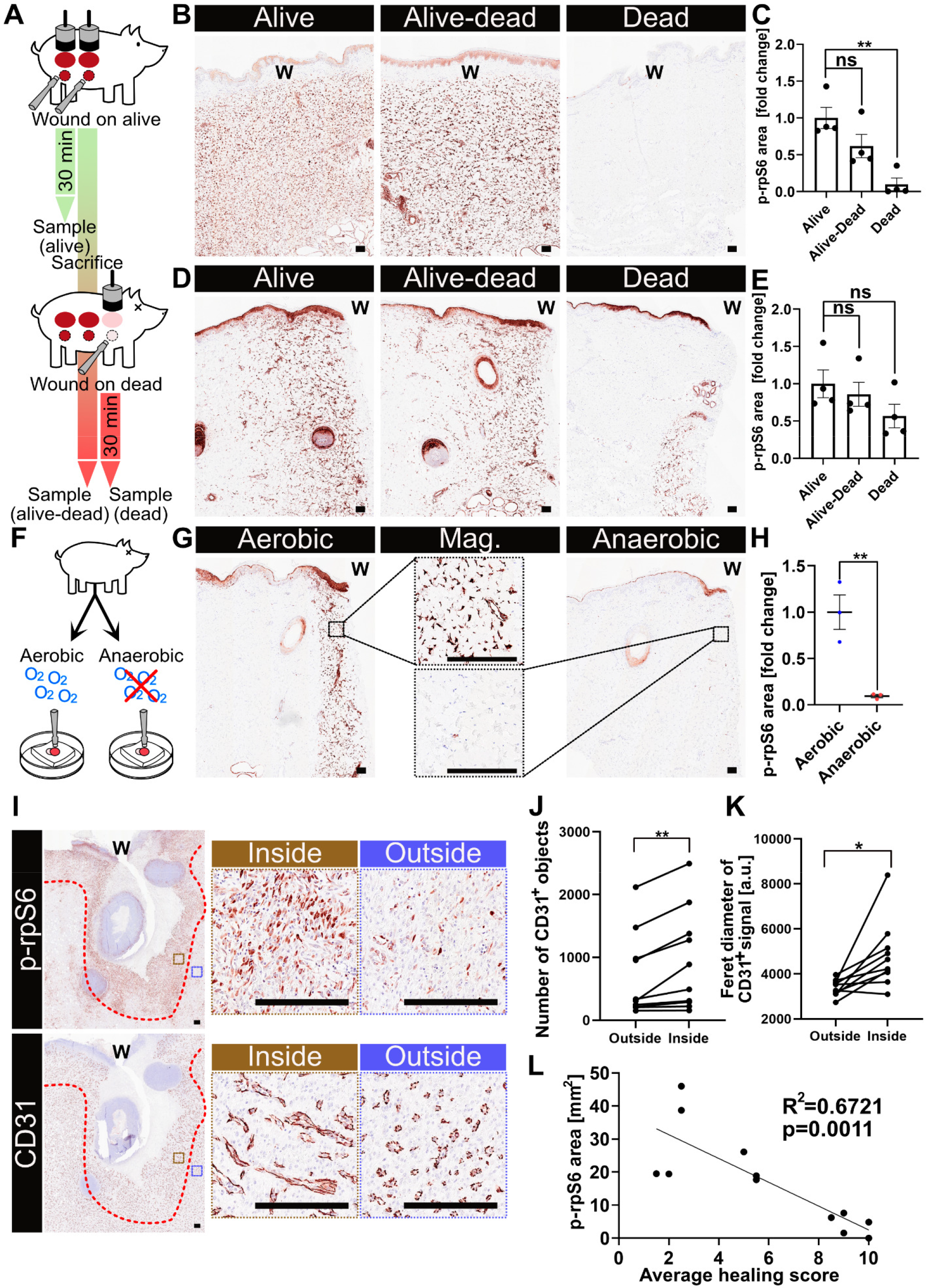
The p-rpS6-zone reports on oxygen availability, vascularization, and progression of healing in pre-clinical studies. (**A**) Experimental model. Pig was injured with two sets of wounds (burn, excision wound), one sample-set was collected after 30 min (condition: “Alive”). The animal was then sacrificed, and a third set of wounds administered. Samples were collected 30 min after death, from injuries performed 60 min prior on the live pig (condition: “Alive-Dead”) and from injuries performed after sacrifice (condition: “Dead”). (**B**) Porcine skin stained for p-rpS6, following burn injury. (**C**) Quantification of p-rpS6 positive area. (**D**) Porcine skin stained for p-rpS6, following excision injury. (**E**) Quantification of p-rpS6 positive area. (**F**) Experimental model. Porcine ex vivo skin was wounded under aerobic (control) or anaerobic conditions. (**G**) Porcine skin stained for p-rpS6, treated as described in panel F. (**H**) Quantification of p-rpS6 positive area. (**I**) Porcine skin samples collected 7 days after wounding and stained for p-rpS6 (upper) and CD31 (lower), with magnifications of regions inside and outside the p-rpS6-zone (marked with a dashed red line). (**J**) Quantification of the number of vessels inside and outside the zone, in equally sized regions. (**K**) Quantification of the Feret diameter of the vessels found inside and outside the zone. (**L**) Correlation between the wound assessment score and the p-rpS6 positive area. Data are from n = 4 pigs for (C) and (E) from n = 3 pigs for (H). For (I)-(M) two samples were collected from each incision wound from 6 pigs at 7 days post injury resulting in n = 12 biological replicates for (M). Out of those, 2 samples, which were fully healed were excluded, thus n =10 biological replicates for the graphs (J) and (K). For all quantifications 10 images per biological replica were used. Mean ± SEM plotted for (C), (E) and (H); for (J) and (K) mean per biological replica is plotted. For the graphs (C), (E) and (H) one-way ANOVA with post-hoc Dunnet’s test was used; for (J) and (K) paired Student’s t-test was used; for (L) Pearson’s correlation test was used. *p<0.05, **p<0.01 and “ns” is “non-significant”. Burn and excision wounds are indicated with a “W”. The scale bars for the images are 100 μm for (b), (d) and (g); for (i) the scale bars show 250 μm for the macro-images, and 100 μm for magnifications.

To verify this hypothesis, we tested the response of ex vivo porcine skin to excision wound injury in conditions deprived of oxygen. To do this, skin was wounded in control (aerobic) conditions or in an anaerobic tent in in the absence of oxygen (Fig. 6 F). As predicted, wounds performed under anaerobic conditions did not develop a p-rpS6-zone (Fig. 6 G and H). In contrast, skin wounded ex vivo under aerobic conditions formed a p-rpS6-zone comparable to that seen under in vivo conditions (Fig. 6 G and H). Interestingly, the zone around the ex vivo excision wound was consistently narrower than the zone seen in wounds performed in vivo, with the depth of the prior reaching approximately 0.3 mm (Fig. 6 G), roughly matching the oxygen penetration depth in tissues (Lovett et al., 2009).

Overall, these results suggest that circulation and oxygenation of tissue are needed for induction, but not maintenance of the p-rpS6-zone. Furthermore, these data show that pathophysiological conditions can exist in which the p-rpS6-zone is disrupted.

### The zone mirrors healing progression

Adequate vasculature is the most important parameter when it comes to the clinical prognosis for the effectiveness of healing (Han and Ceilley, 2017). As the p-rpS6-zone is functionally linked to tissue oxygenation we considered the possibility of using the zone to characterize healing states of wounds. To challenge the diagnostic value of an assessment of the formation of the p-rpS6-zone in a pre-clinical scenario, we characterized the properties of the zone in porcine skin samples collected 7 days post-wounding.

Pigs underwent a surgery unrelated to this study, which required a central laparotomy incision into the abdominal cavity and was closed with resorbable sutures. At sacrifice, 7 days post-surgery, differences in the healing process due to inter-animal variation were observed, even within the same injury (Fig. S7 C). Two trained veterinarians provided an unbiased wound score for each position of the wound where a sample was taken, where 1 represents an unhealed wound and 10 represents a fully healed wound (Fig. S7 C).

In order to assess the relationship between vasculature and the p-rpS6-zone we quantified the number and size of blood vessels inside the zone and a size-matched region just outside of the zone in the same sample (Fig. 6 I). We found that the tissue within the zone consistently demonstrated a larger quantity of blood vessels, which were also of a larger average size (Fig. 6 I to K). It is worth stressing that this property of the zone is so consistent it enables the clear discrimination between regions of poor and rich vasculature.

We correlated the wound score with the area occupied by the p-rpS6-zone for each sample and found that the two are strongly inversely correlated: wounds with a larger p-rpS6-zone show a lower score and appear less healed (Fig. 6 L and fig. S7 D, E). We also found that even in wounds that macroscopically appear healed, there is a small quantity of p-rpS6 left, that could be used to trace where the wound used to be (Fig. S7 F). In summary, these results demonstrate that the p-rpS6-zone accurately reports on the status of the skin vasculature and healing efficiency in pre-clinical samples and thus can be used as a measure of healing status.

## Discussion

The temporal dynamics of tissue response to injury and the healing process is relatively well understood with methods such as real-time PCR and western blotting providing essential information on the levels of factors driving inflammation, matrix remodeling and angiogenesis (Rodrigues *et al*., 2019). However, the spatial characterization of these processes is still largely unknown with only recent research work suggesting how tissues of non-mammalian model organisms govern generation and propagation of signals necessary for healing (De Simone *et al*., 2021; Toyota *et al*., 2018). Multiple questions remain unanswered, such as how deep inside the tissue will cells undergo cell death and which will survive; where in the tissue does the response to wounding start and end, and what part of the damaged tissue initiates and drives its healing?

Here, we show a phenotype of spatial response of skin to damage in mammals. The p-rpS6-zone defines an area of skin tissue starting at the end of the necrotic layer at the side of the skin injury. The zone forms within minutes after damage and continues throughout the process of healing. One of the features of the p-rpS6-zone is its stability and ease of detection. While other spatially-organized markers that originate from wounding such as ROS gradient (Niethammer et al., 2009) or p-Erk waves (De Simone *et al*., 2021) operate on a basis of propagation of signal spikes or gradients that spread across the tissue, the p-rpS6-zone establishes itself as a stable zone of signal around the insult. This makes it easily detectable in all laboratories without usage of transgenic animals or in vivo reporter systems.

Despite the wide range of phenotypes characterizing response to wounding and the healing process, the one marker capable of defining a general response of skin to the presence of a wound is still missing. The p-rpS6-zone appears to comprise the core cellular processes related to healing, including cell growth, proliferation, senescence and angiogenesis. Although p-rpS6 has been seen in wounds before, it has never before been characterized as a stand-alone feature (Squarize et al., 2010; Tsai et al., 2022). The definition of the p-rpS6-zone brings with it several advantages over classic histological wound size assessment and extrapolation of its spatial properties onto the surrounding skin architecture. For one, it is valuable to have an easily detectable single marker that represents multiple other features of healing and delineates the tissue which has responded to the initial damage. Also, it is likely that the p-rpS6-zone accurately reports on healing defects such as poor vasculature and hypoxia. Finally, the p-rpS6-zone is consistently activated by a wide range of wound types, including burns, excision, incision or needle pricks, which to our knowledge is unique. Thus, the characterization of the zone provides clinically relevant data that goes beyond the identification of the wounded region.

Causes and consequences of the p-rpS6-zone formation, however, remain partially unresolved. For the prior, we could show that the zone is a consequence of the contact between cells and extracellular DAMPs that lead to rpS6 phosphorylation through an mTOR-dependent mechanism. It remains to be determined whether any specific subset of DAMPs is sufficient to induce zone formation and whether other mTOR-interacting proteins respond in a similar manner. In fact, it is possible that the upstream regulators differ between cell types as varied as endothelial cells and keratinocytes, and could include such diverse players as calcium influx, and kinases (PKC, PKA and RSK). This only strengthens the value of p-rpS6 as a marker of wounding, as it seems to represent a global tissue injury response of all cell types.

As for the consequences of the zone, our experiments in phospho-deficient transgenic mice reveal a possible role of the p-rpS6-zone in slowing the initial tissue response by increasing the expression of a senescent phenotype in keratinocytes and reducing their proliferation. In contrast, the p-rpS6-zone appears to be beneficial for the final resolution of a wound, as the phospho-deficient mouse model showed disrupted wound healing with increased αSMA deposition and an abnormal collagen I to collagen III ratio. This could further drive the hypothesis that an early induction of senescence upon wounding is required for effective healing and regeneration (Nishiguchi et al., 2018).

A key observation in this research work is that the spherical structure of the p-rpS6-zone which encapsulates the wound and is characterized by increased proliferation, migration, and angiogenesis, mirrors many features of a malignant tumor. Indeed, cancer has been characterized as a “wound that never heals” (Deyell et al., 2021), and the phosphorylation of rpS6 itself is a feature frequently found in tumors (Yi et al., 2021). This may indicate that a wound could in fact be a “cancer that heals”. Future research will have to determine if the wound-induced p-rpS6-zone constitutes a cancer-like structure.

In conclusion, the p-rpS6-zone occurs in animal skin as an immediate response of cells affected but not killed by an injury. The zone is unique as it divides an otherwise homogenous tissue, visually marking an area of injury-response, separate from homeostatic tissue. In pre-clinical samples the zone reports on vascularization and is predictive of the extent of healing. The p-rpS6-zone promises to be an invaluable clinical and diagnostic tool.

## Acknowledgments

The Research Group Senescence and Healing of Wounds (SHoW) is a collaboration between the Ludwig Boltzmann Gesellschaft GmbH and the Austrian Workers’ Compensation Board (AUVA) with support of the Austrian Nationalstiftung. We would like to thank Peter Dungel and Magdalena Metzger for their help with mouse experiments. We would like to thank the LBI Trauma experimental surgery team including Astrid Hönigsberger, Karin Brenner and Elmar Ebner, as well as Roberto Plasenzotti and Rebecca Nistelberger from the Medical University of Vienna for help with animal experiments and husbandry. We would like to thank Anja Wagner for help with protein isolation and assistance in other experiments. We would like to thank Matthias Horn and Angelika Schwarzhans from the Centre for Microbiology and Environmental Systems Science, Vienna for access to their anoxia chamber. We would like to thank Mario Pende for sharing the rpS6 KI mice. We would like to acknowledge our funding sources: der Wissenschaftsfonds grant P 35382 (M.O., B.B., T.R.), der Wissenschaftsfonds grant P 35268-B (J.G.), and the Lorenz Böhler Gesellschaft grant LBF 6/21 (B.B.).

## Author contributions

Conceptualization: MO, NARR, HD

Methodology: NARR, HD, BB, BS, KV, TR, PH, KS, GL, JF, SD, MM, MS, PS, OM, FG

Investigation: MO, NARR, HD, BB, BS, KV, TR, PH

Supervision: MO

Writing – original draft: MO

Writing – review & editing: MO, NARR, JG, HR

## References

Brugarolas, J., Lei, K., Hurley, R.L., Manning, B.D., Reiling, J.H., Hafen, E., Witters, L.A., Ellisen, L.W., and Kaelin, W.G., Jr. (2004). Regulation of mTOR function in response to hypoxia by REDD1 and the TSC1/TSC2 tumor suppressor complex. Genes Dev 18, 2893–2904. 10.1101/gad.1256804.

Celli, A., Tu, C.L., Lee, E., Bikle, D.D., and Mauro, T.M. (2021). Decreased Calcium-Sensing Receptor Expression Controls Calcium Signaling and Cell-To-Cell Adhesion Defects in Aged Skin. J Invest Dermatol 141, 2577–2586. 10.1016/j.jid.2021.03.025.

Cordeiro, J.V., and Jacinto, A. (2013). The role of transcription-independent damage signals in the initiation of epithelial wound healing. Nat Rev Mol Cell Biol 14, 249–262.

Cox, J. (2013). Pressure ulcer development and vasopressor agents in adult critical care patients: a literature review. Ostomy Wound Manage 59, 50-54, 56-60.

De Simone, A., Evanitsky, M.N., Hayden, L., Cox, B.D., Wang, J., Tornini, V.A., Ou, J., Chao, A., Poss, K.D., and Di Talia, S. (2021). Control of osteoblast regeneration by a train of Erk activity waves. Nature 590, 129–133. 10.1038/s41586-020-03085-8.

Demaria, M., Ohtani, N., Youssef, S.A., Rodier, F., Toussaint, W., Mitchell, J.R., Laberge, R.M., Vijg, J., Van Steeg, H., Dolle, M.E., et al. (2014). An essential role for senescent cells in optimal wound healing through secretion of PDGF-AA. Dev Cell 31, 722–733. 10.1016/j.devcel.2014.11.012.

Deyell, M., Garris, C.S., and Laughney, A.M. (2021). Cancer metastasis as a non-healing wound. Br J Cancer 124, 1491–1502. 10.1038/s41416-021-01309-w.

Han, G., and Ceilley, R. (2017). Chronic Wound Healing: A Review of Current Management and Treatments. Adv Ther 34, 599–610. 10.1007/s12325-017-0478-y.

Hewitt, G., Jurk, D., Marques, F.D., Correia-Melo, C., Hardy, T., Gackowska, A., Anderson, R., Taschuk, M., Mann, J., and Passos, J.F. (2012). Telomeres are favoured targets of a persistent DNA damage response in ageing and stress-induced senescence. Nat Commun 3, 708. 10.1038/ncomms1708.

Holzer, J.C.J., Tiffner, K., Kainz, S., Reisenegger, P., Bernardelli de Mattos, I., Funk, M., Lemarchand, T., Laaff, H., Bal, A., Birngruber, T., et al. (2020). A novel human ex-vivo burn model and the local cooling effect of a bacterial nanocellulose-based wound dressing. Burns 46, 1924–1932. 10.1016/j.burns.2020.06.024.

Lovett, M., Lee, K., Edwards, A., and Kaplan, D.L. (2009). Vascularization strategies for tissue engineering. Tissue Eng Part B Rev 15, 353–370. 10.1089/ten.TEB.2009.0085.

Martin, P., and Nobes, C.D. (1992). An early molecular component of the wound healing response in rat embryos--induction of c-fos protein in cells at the epidermal wound margin. Mech Dev 38, 209–215. 10.1016/0925-4773(92)90054-n.

Meyuhas, O. (2015). Ribosomal Protein S6 Phosphorylation: Four Decades of Research. Int Rev Cell Mol Biol 320, 41–73. 10.1016/bs.ircmb.2015.07.006.

Moreira, S., Stramer, B., Evans, I., Wood, W., and Martin, P. (2010). Prioritization of competing damage and developmental signals by migrating macrophages in the Drosophila embryo. Curr Biol 20, 464–470. 10.1016/j.cub.2010.01.047.

Niethammer, P., Grabher, C., Look, A.T., and Mitchison, T.J. (2009). A tissue-scale gradient of hydrogen peroxide mediates rapid wound detection in zebrafish. Nature 459, 996–999. 10.1038/nature08119.

Nishiguchi, M.A., Spencer, C.A., Leung, D.H., and Leung, T.H. (2018). Aging Suppresses Skin-Derived Circulating SDF1 to Promote Full-Thickness Tissue Regeneration. Cell Rep 24, 3383–3392 e3385. 10.1016/j.celrep.2018.08.054.

Ogrodnik, M., Salmonowicz, H., Jurk, D., and Passos, J.F. (2019). Expansion and Cell-Cycle Arrest: Common Denominators of Cellular Senescence. Trends Biochem Sci 44, 996–1008. 10.1016/j.tibs.2019.06.011.

Rocha, S.F., Schiller, M., Jing, D., Li, H., Butz, S., Vestweber, D., Biljes, D., Drexler, H.C., Nieminen-Kelha, M., Vajkoczy, P., et al. (2014). Esm1 modulates endothelial tip cell behavior and vascular permeability by enhancing VEGF bioavailability. Circ Res 115, 581–590. 10.1161/CIRCRESAHA.115.304718.

Rodrigues, M., Kosaric, N., Bonham, C.A., and Gurtner, G.C. (2019). Wound Healing: A Cellular Perspective. Physiol Rev 99, 665–706. 10.1152/physrev.00067.2017.

Ruvinsky, I., Sharon, N., Lerer, T., Cohen, H., Stolovich-Rain, M., Nir, T., Dor, Y., Zisman, P., and Meyuhas, O. (2005). Ribosomal protein S6 phosphorylation is a determinant of cell size and glucose homeostasis. Genes Dev 19, 2199–2211. 10.1101/gad.351605.

Schindelin, J., Arganda-Carreras, I., Frise, E., Kaynig, V., Longair, M., Pietzsch, T., Preibisch, S., Rueden, C., Saalfeld, S., Schmid, B., et al. (2012). Fiji: an open-source platform for biological-image analysis. Nat Methods 9, 676–682. 10.1038/nmeth.2019.

Squarize, C.H., Castilho, R.M., Bugge, T.H., and Gutkind, J.S. (2010). Accelerated wound healing by mTOR activation in genetically defined mouse models. PLoS One 5, e10643. 10.1371/journal.pone.0010643.

Sullivan, T.P., Eaglstein, W.H., Davis, S.C., and Mertz, P. (2001). The pig as a model for human wound healing. Wound Repair Regen 9, 66–76. 10.1046/j.1524-475x.2001.00066.x.

Toyota, M., Spencer, D., Sawai-Toyota, S., Jiaqi, W., Zhang, T., Koo, A.J., Howe, G.A., and Gilroy, S. (2018). Glutamate triggers long-distance, calcium-based plant defense signaling. Science 361, 1112–1115. 10.1126/science.aat7744.

Tsai, C.L., Changchien, C.Y., Chen, Y., Chang, H.H., Tsai, W.C., Wang, Y.W., Chou, K.C., Chiang, M.H., Tsai, Y.L., Tsai, H.C., et al. (2022). Accelerated Wound Healing and Keratinocyte Proliferation through PI3K/Akt/pS6 and VEGFR2 Signaling by Topical Use of Pleural Fluid. Cells 11. 10.3390/cells11050817.

van Vliet, T., Varela-Eirin, M., Wang, B., Borghesan, M., Brandenburg, S.M., Franzin, R., Evangelou, K., Seelen, M., Gorgoulis, V., and Demaria, M. (2021). Physiological hypoxia restrains the senescence-associated secretory phenotype via AMPK-mediated mTOR suppression. Mol Cell 81, 2041–2052 e2046. 10.1016/j.molcel.2021.03.018.

Yi, Y.W., You, K.S., Park, J.S., Lee, S.G., and Seong, Y.S. (2021). Ribosomal Protein S6: A Potential Therapeutic Target against Cancer? Int J Mol Sci 23. 10.3390/ijms23010048.

Zhao, R., Liang, H., Clarke, E., Jackson, C., and Xue, M. (2016). Inflammation in Chronic Wounds. Int J Mol Sci 17. 10.3390/ijms17122085.

