## Supplemental data for "The p-rpS6-zone delineates wounding response and the healing process"

### Materials and Methods

Ex vivo and in vivo wounding experiments in pigs

Pig experiments were performed according to the ethical guidelines of the host institution with support of the veterinarian team of the LBI Trauma. All experimental protocols were approved in advance by the Municipal Government of Vienna in accordance with Austrian law and the Guide for the Care and Use of Laboratory Animals as defined by the National Institute of Health (revised 2011). These protocols allow for non-recovery experiments with biopsy and sample collection up to 6 h post injury. Samples were collected from a total of 26 male pigs that were also being used for other experiments.

Pigs were anesthetized using zoletil (250 mg of tiletamine with 250 mg of zolazepam) mixed with 100 mg of xylazine in 5 ml of solution – of this 1-2 ml were administered intramuscularly per 15 kg of body weight. A catheter was then placed in the lateral ear vein and 2 mg/kg propofol (10%) were administered intravenously. Pigs were then intubated and ventilated. During the procedure pigs were kept under inhalation anesthesia (oxygen, sevoflurane 3-6%). A pulse oximeter (SpO2, heart rate), and temperature probe (esophageal) were additionally used for monitoring animals. In addition, the pigs received intravenous 2-10 ml/kg bw Elomel isoton® (Fresenius Kabi, Graz, Austria), 0.008 mg/kg/h sufentanil, and 2.5 mg/kg/h rocuronium bromide while under anesthesia. During the surgical procedure, the sufentanil dose was adjusted according to the condition of the animals and close attention was paid to provide adequate analgesia. If blood pressure dropped, 5 µg/kg/h L-norepinephrine hydrochloride was administered intravenously until effect (MAP 60>mmHg). Operation logging was performed every 15 min. Further instrumentation of the animals consisted of preparation of the carotid artery (monitoring of central arterial blood pressure) and jugular vein (infusion, administration of analgesic and muscle relaxant) and placement of a bladder catheter.

Subsequently, wounds were applied to the shaved dorsal skin. Wounds were induced by burn (standardized thermal lesions of grade I or grade IIa), biopsy punch (as a proxy for excision wounds) or needle prick (29 G U-40 insulin needle). For burns, an aluminum block with a diameter of 3 cm on a specially designed plunger was heated to 60, 70 or 80 °C in a water bath, the temperature of the surface of the block was measured using an infrared contactless thermometer, and the plunger was pressed with a constant and normalized across experiments pressure of approximately 0.4 $kg/cm^2$ onto the side of the pig. Samples were collected at specified time points after the burn, specifically 5, 15 and 30 min and 1.5, 3 and 6 h. Excision wounds were modelled by performing a 6 mm biopsy punch of the skin, removing the biopsy and leaving a 6 mm diameter wound. The edges of this wound were then collected at the same time points as for the burn model. Samples were immediately fixed in 10% formalin for 24 h at room temperature in histology cassettes. They were then washed carefully for 1 h with water, incubated for 1 h in 50% ethanol and stored in 70% ethanol until the samples were finally embedded in paraffin.

For the experiments focused on hypoxia, two sets of burn and excision wounds were performed on the dorsal skin of an alive and anesthetized pig. The first set of samples was collected after 30 min, after which the pig was sacrificed. The second set of samples was collected 30 min after death. Before that, approximately 3 min after death, a third set of burn and excision wound was placed near the first two, and samples were collected after 30 min. For this set of experiments, temperature measurements of the skin were taken using an infrared temperature gun immediately prior to the burn, the temperature of the burned skin was measured immediately after the burn and again at sample collection to control for fluctuations in body temperature caused by death.

For ex vivo experiments, dorsal porcine skin was removed from the sacrificed animal and kept for acclimatization for 1 h in an incubator at 37 °C. The tissue was then subjected to wounding using a 6 mm biopsy punch. At specific time points the edge of the original biopsy-wound was extracted and fixed in formalin. To analyze the effect of wounding in anaerobic conditions, ex vivo skin was transported in a warmed box into an anaerobic tent. One piece of skin was wounded and left to recover in an incubator at 37 °C in atmospheric oxygen, while the second piece was placed in an anaerobic tent for wounding and then left to recover at 37 °C remaining under anaerobic conditions.

In vivo wounding experiments in BALB/c mice

For the experiments in wild type mice as shown in the main figures 1 and 3, and extended figures 1 and 4 wild-type (WT), female BALB/c mice were purchased from Janvier labs. Mouse experiments were performed according to ethical guidelines of the host institution with support of the veterinarian team of the LBI Trauma. All experimental protocols were approved in advance by the Municipal Government of Vienna in accordance with Austrian law and the Guide for the Care and Use of Laboratory Animals as defined by the National Institute of Health (revised 2011). 20 mice were used for experiments at an age of 12 weeks and an average weight of 20–25 g. Mice were wounded and then sacrificed at 0.5-1.5 h, 3 d, 7 d, 12 d and 28 d after wounding.

For all experiments, a maximum of five animals were housed per Type-III cage on a 12 h light–dark diurnal cycle with room temperature between 21 and 23 °C. Standard rodent diet (Abbedd Lab & Vet Service, Vienna, Austria) and water were provided prior to and throughout the experiments. The excision wounds were performed under inhalation anesthesia using an oxygen/sevoflurane mixture (4-5 %) (Sevorane®, AbbVie Inc., North Chicago, Illinois, USA). Analgesia (1 mg/kg of meloxicam p.o.) was administered 120 min prior to surgery, as well as daily until the end of the study or until wound closure.

All mice were wounded using the following wounding protocol. Mice were anesthetized, the hair was shaved in the dorsal area and the skin was disinfected with Isozid®. A template was used to draw a 1 cm diameter circle on the posterior dorsal region and the skin was then carefully cut along this circle using scissors to avoid damaging the underlying muscle fascia, creating a single full-thickness excision wound. After wounding, mice were placed in a cage on a 37 °C heating pad until recovery. Then the mice were transferred back to their cages and housed for the duration of the study. Animals were monitored daily for any signs of infection, pain and loss of weight. To follow the healing process, digital photos were taken using a camera (LifeViz micro, Quantificare, France). Wound area was analyzed by a planimetric measurement with a free-hand tool using Fiji image processing software (Schindelin et al., 2012) (ImageJ, National Institute of Health, USA). Mice were sacrificed at the end of the observation period under deep sevoflurane anesthesia via cervical dislocation. Each wound was collected with the surrounding tissue and placed in a histology cassette. The tissues were fixed in 10% formalin for paraffin sections.

In vivo wounding experiments in C57BL/6 and rpS6 p-/- knock-in mice

For experiments on C57BL/6 wild type (WT) mice and transgenic rpS6 phospho-deficient knock-in mice rpS6 p-/- (rpS6 KI) mice which have a C57BL/6 background, all experimental protocols were approved in advance by the Austrian Bundesministerium für Bildung, Wissenschaft und Forschung (BMBWF). RpS6 KI mice were compared to age- and sex-matched C57BL/6 WT mice bred in the same facility, at an average age of 11 weeks. All mice were housed and wounded as described above, except for a slight modification to the analgesia protocol. Analgesia was administered at least 30 min prior to surgery (6 mg/kg carprofen p.o.) and post-operatively in their drinking water for three days.

Healing kinetics were performed in females and males: eight female rpS6 KI mice and ten age-matched female WT mice bred in the same facility, and eight male rpS6 KI mice and nine age-matched male WT littermates (rpS6$^{+/+}$). All mice were wounded as described above and sacrificed at 13 days post-wounding. A further cohort of mice was sacrificed at 3 days post-wounding and consisted of six male rpS6 KI mice and seven male WT mice.

Wounds were photographed using a digital camera (LifeViz micro, Quantificare, France) and quantified as described above, by a planimetric measurement with a free-hand tool using Fiji image processing software (Schindelin *et al.*, 2012) (ImageJ, National Institute of Health, USA). The remaining unhealed wounds were quantified using the same methodology, taking into consideration the entire hair-less and freshly formed epidermis surrounding the remaining wound bed.

*Ex vivo* experiments in human skin

The skin samples for the sections used in this study were approved by the Ethics Committee of the Medical University of Vienna (1969/2021), and written informed consent was obtained from all subjects. Tissue was used for experiments within 5 hours from removal, throughout which the tissue was stored at room temperature. Prior to the experiment the tissue was acclimatized in an incubator at 37 °C and wounded using the same procedure applied to ex vivo porcine skin.

Histology

For sample analysis both immunohistochemistry and immunohistofluorescence were used. Skin samples were fixed in 10 % buffered formalin for 24 h and, embedded in paraffin and sectioned from the center of the wound at a thickness of 4 µm. Sections were deparaffinized and rehydrated through a graded alcohol series. For heat-induced antigen retrieval samples were incubated in a 0.1 M Tris with 0.01 M EDTA (pH 9.0) or sodium- citrate buffer 0.01 M (pH 6.0) in a steamer at 95°C for 20 min, or in the microwave where the buffer was brought to a boil and then left for 15 min at sub-boiling temperature.

For immunohistofluorescence, samples were blocked in PBS containing 0.4 % BSA (Sigma Aldrich) and 1.6 % normal goat serum (Vectorlabs) for 1 h at room temperature. Samples were then incubated with appropriate primary antibodies (Table 1) in blocking buffer overnight at 4 °C. Slides were washed three times with TBS-Tween (TBS-T) and incubated for 1 h with DAPI and fluorescent secondary antibody (ThermoFisher) and mounted in MOWIOL mounting media.

For immunohistochemistry, samples were blocked for 10 min using BLOXALL (Vectorlabs), were washed in TBS-T and were then incubated for 1 h at room temperature overnight at 4°C with primary antibody (Table 1). After incubation, samples were washed three times, incubated with HRP-conjugated secondary antibody (BrightVision) for 30 min and stained for 6 min with NovaRed (Vectorlabs). Finally, samples were counterstained, dehydrated and mounted. All applied primary antibodies can be seen in Table 1 and were used in dilutions recommended by the manufacturer.

To analyze immunohistochemistry stainings, the stained area was quantified by measuring the amount of the bound antibody using ImageJ. Specifically, slide-scans were exported to the tif format, and the number of positive pixels was quantified in experiment-specific regions of interest (ROI) using "Color deconvolution" and vectors for "H DAB" followed by signal thresholding and "Analyze Particles" function to determine the stained area. All the settings were consistent for the samples within a given experiment.

The analysis of frequency and size of CD31-positive objects in relation to p-rpS6 for porcine and human samples was done by first aligning histological images in Fiji (Schindelin *et al.*, 2012) using the moving least squares plugin. Between 10 and 20 points which could be identified on both images were marked using the multi point selection tool. The CD31 stained image was warped to match the ps6 staining. Due to the size of the images, the RGB color channels were split into three separate images, warped individually using the moving least squares plugin with the "rigid" method and then merged again into a single RGB image. Alignment errors were corrected by moving or adding points and repeating the warping procedure. The aligned images were then processed as described above. P-rpS6-positive area was selected and the ROI was copied onto the CD31-stained section. In this way two images of CD31 staining were obtained, one of the inside of the p-rpS56 zone and a second for which the same ROI was moved outside of the p-rpS6-zone (thus the selected areas are identical in shape and size). Both images were colour-deconvolved, thresholded for positive signal and analyzed using "Particle analysis" tool in ImageJ deriving the number ("Count") and size ("Feret") of the CD31-positive objects inside and outside of the zone.

The histological analysis based on the assessment of depth of the initiation of the signal was performed in OlyVIA software (Olympus Corporation, Tokyo, Japan). For this analysis several measurements of the length between epidermis and the positive area were made and the average compared between animals and conditions.

Depth Probability Chart

For each timepoint after burn or excision wound, mean and standard deviation were calculated using Google Sheets (Google LLC). The depths were assumed to follow a normal distribution and were linearly interpolated between measured timepoints using the "Forecast. Linear" function. Probabilities for each timepoint and depth were calculated using the cumulative "Normdist" function. Confidence interval for each timepoint was calculated using the "Confidence" function. To convert the calculated probabilities into an image, probabilities for each timepoint and depth were exported as a tab separated text file. Using Fiji, an empty 32-bit image was created and each pixel assigned the probability value in the tab separated text file using a macro. Color values were assigned to each probability using a lookup table.

Needle 3D zone image and its cross-sectional view

To create the 3D segmentation and virtual cross section, serial histological sections were stained. The sample was sectioned horizontally starting from the epidermis into 40 slices of 4 µm thickness with 20 µm gaps between each slice. All 40 slices were stained for p-rpS6, acquired on a scanning microscope and then used to create a 3D projection representing regions of tissue positive for p-rpS6 staining. Images were aligned using Fiji (Schindelin *et al.*, 2012). First, distortions of the individual histological slides were corrected by creating point ROIs which marked the position of the same hair follicles on each slide. Using the moving least squares plugin, the slides were distorted so the position of the hair follicles was identical for all slides. This procedure removes distortions from the cutting and mounting of the slides, but since not all follicles run parallel, some misalignment remains. To correct this, images were rigidly registered by drawing a line between two follicles which lie on opposing sides of the defect and are approximately equidistant. The line was then moved so the center of the line lies in the center of the defect. Images were then rotated so the angle of the lines for all slides matches the line on the first slide and translated so the center of the line is in the same position for all slides. Registered images were combined into a single stack.

The outline of the affected area was marked using the lasso tool and saved with the ROI manager. Segmentations for the skin and affected area were saved as binary images. 3D rendering was created using CTVox (Bruker Corporation, Billerica, MA, USA). To create the virtual cross section, a rectangular selection with a height that matches the slice distance was placed over the center of the defect. The stack was then cropped to this selection and a montage was created where all slices in the cropped stack are arranged in a single column.

*In vitro* experiments

To model a skin wound in vitro, cell cultures were subjected to a damage assay to monitor cellular response mechanisms. Commercially available human dermal fibroblasts (HDFs) were cultured in HDF medium (DMEM/F12 (Sigma Aldrich), 10 % fetal bovine serum (Sigma Aldrich) and 4 mM L-Glutamine (Gibco)), subcultivated at 90 % confluency and used for experiments below passage 15. Immortalized human keratinocytes (HaCaT) were maintained in HACAT medium (DMEM high glucose (Sigma Aldrich), 10 % fetal bovine serum (Sigma Aldrich) and 2 mM L-Glutamine (Gibco) and subcultivated at 60 % confluency. All cell culture experiments were performed in a humidified incubator at 37 °C and 5 % CO2.

The scratch assays were performed on cells seeded at 7,000 cells/cm2 (HDF) 10,000 cells/cm2 (HACAT) and grown for 5 to 7 days until confluency. 24 h prior to scratching, medium was exchanged to basal medium (DMEM/F12 or DMEM without any supplements) to eliminate endogenous activation of signaling pathways. For signaling pathway manipulation, cells were treated using either vehicle (0.1 % DMSO), 20 nM of mTOR inhibitor rapamycin, or 10 µM of MEK inhibitor U0126 for 10 to 30 min prior to scratch or 30 min after administration of the media with DAMPs. For the scratch assay, the confluent cell layers were then subjected to damage by scraping a blunt-edged 21 g needle through the confluent cell layer, thereby creating a standardized scratch with broken cells on either edge. All scratches were verified using an inverted microscope directly after scratch. After 5, 30 or 90 min of incubation the cell layers were fixed using 10 % formalin for 10 min, washed, and stored at 4 °C until they were stained.

For immunofluorescence staining, the coverslips containing damaged and control cells were permeabilized using 0.5 % Triton X-100 for 5 min, washed three times with PBS for 5 min each, and blocked using PBS with 5 % goat serum (VECS-1000, Vector Laboratories) for 1 h. Subsequently, primary antibody and Phalloidin-iFluor 488 (Abcam, ab176753) were applied in optimized dilution and incubated overnight at 4 °C. After primary antibody application, the coverslips were washed three times for 5 min and secondary antibody and DAPI were applied for 1 h, cells were washed, and mounted on slides using MOWIOL solution. For quantification of rpS6 and Erk phosphorylation, ten images per slide were taken along the scratch using an inverted microscope Nikon (Nikon Corporation, Tokyo, Japan) with set exposure times. These images were processed and quantified for total cell count and activated cell count (p-rpS6 or p-ERK positive cells) using set thresholds and particle analysis in Fiji. All applied primary antibodies were used in dilutions recommended by the manufacturer: αSMA (A2547, Sigma), CD3 (MCA1477, Biorad), cFos (ab208942, Abcam), Cleaved Caspase 3 (9661, Cell Signaling), E-Cadherin (M3612, Dako), γ-H2A.X (9718, Cell Signaling), Hmgb1 (ab18256, Abcam), p-Erk (Thr202/Tyr204) (4370, Cell Signaling), p-rpS6 (S235 + S236) (4858, Cell Signaling), p-rpS6 (S240 + S244) (ab81081, Abcam), p21 (ab107099, Abcam), PCNA (ab29, Abcam), total rpS6 (2317, Cell Signaling), Endocan (AF1999, Novus Biologicals), CD31 for IHC (sc-376764, Santa Cruz), CD31 for IHF (DIA-310, Dianova), K10 (GP-K10, Progen), K16 (GP-K16, Progen).

Western blotting

Skin biopsies were snap frozen in liquid nitrogen until processing. Tissue was then pulverized using a mortar and pestle while submerged under liquid nitrogen. Tissue powder was lysed using RIPA lysis buffer (Sigma-Aldrich, 20-188) supplemented with protease and phosphatase inhibitors (Sigma-Aldrich, P8340 and P0044), 0.2 % SDS and 0.5 mM DTT. Samples were lysed for 1 h on ice, then sonicated 3 times at 0.5 kJ per pulse and finally centrifuged 4 times at 12,000 xg for 10 min to remove all fat from the lysate. Samples quantified using the Pierce BCA protein assay kit, boiled in NuPAGE LDS Sample Buffer (ThermoFisher) and run on Bolt 10 % Bis-Tris Plus Gels, 12-well gels (ThermoFisher) in MOPS buffer. The protein was transferred to a PVDF membrane using the Bradford Trans-blot Turbo Transfer system for protein detection. Membranes were incubated in primary antibody overnight (p-rpS6 (S240 + S244), ab81081, Abcam; total rpS6, 2217, Cell Signaling) then incubated with secondary HRP-conjugated antibody for 1 h and developed using ECL (ThermoFisher). GAPDH was used as a normalization control (Gapdh, MA5-15738, ThermoFisher).

Statistical Analysis

All statistical analyses including testing the normality of data distribution were performed using GraphPad Prism 9.3.1 and a P value <0.05 was considered as significant. For differences between 2 groups paired or unpaired two-tailed t-test was used, data were further tested for equality of variances using F test. For >2 group comparisons, one-way ANOVA with Dunnet's multiple comparison test was used. For analysis concerning more than one variable two-way ANOVA with

Sidak’s multiple comparison test or multiple t test was used. Correlations were assessed using Pearson’s rank correlation test.

### Supplemental figures and legends

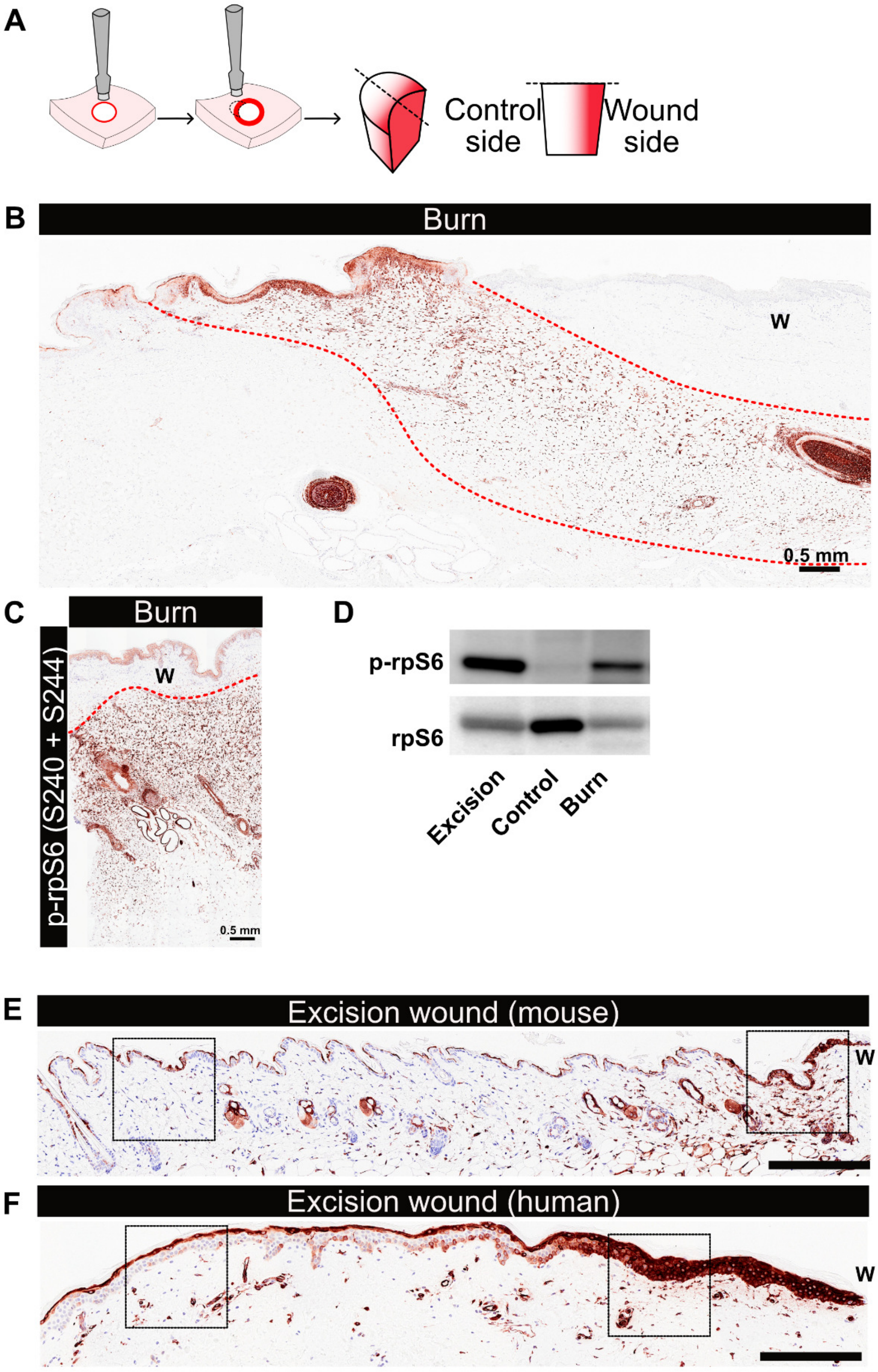

**Fig. S1. RpS6 phosphorylation marks a zone of tissue activation in response to wounding, a damage response which is conserved in mammal.**

(**A**) A scheme demonstrating the use of a biopsy punch as a proxy for excision wound, and the subsequent sample collection. (**B**) An image of porcine skin biopsy collected from a side of a 60 ˚C burn injury representing p-rpS6 IHC staining. (**C**) Porcine skin stained with an antibody recognizing the alternative phosphorylation site of p-rpS6 (S240/S244) in 60 ˚C burn injury sample. (**D**) Western blot for p-rpS6 (S235/236) and GAPDH (loading control) in porcine skin samples collected 1.5 h after excision injury, 60 ˚C burn injury or control. (**E**) An image of mouse skin showing p-rpS6 staining. Frames mark regions for which micrographs were taken (for the Figure 1) to show p-rpS6 staining in regions proximal (Excision site) and distal (Control site) to the wound site (the right side of the image). (**F**) An image of human ex vivo skin showing p-rpS6 staining. Frames mark regions for which micrographs were taken (for the Figure 1) to show p-rpS6 staining in regions proximal (excision site) and distal (control site) to the wound (the right side of the image). The scale bars for (B) and (C) show 500 µm and for (E) and (F) show 100 µm.

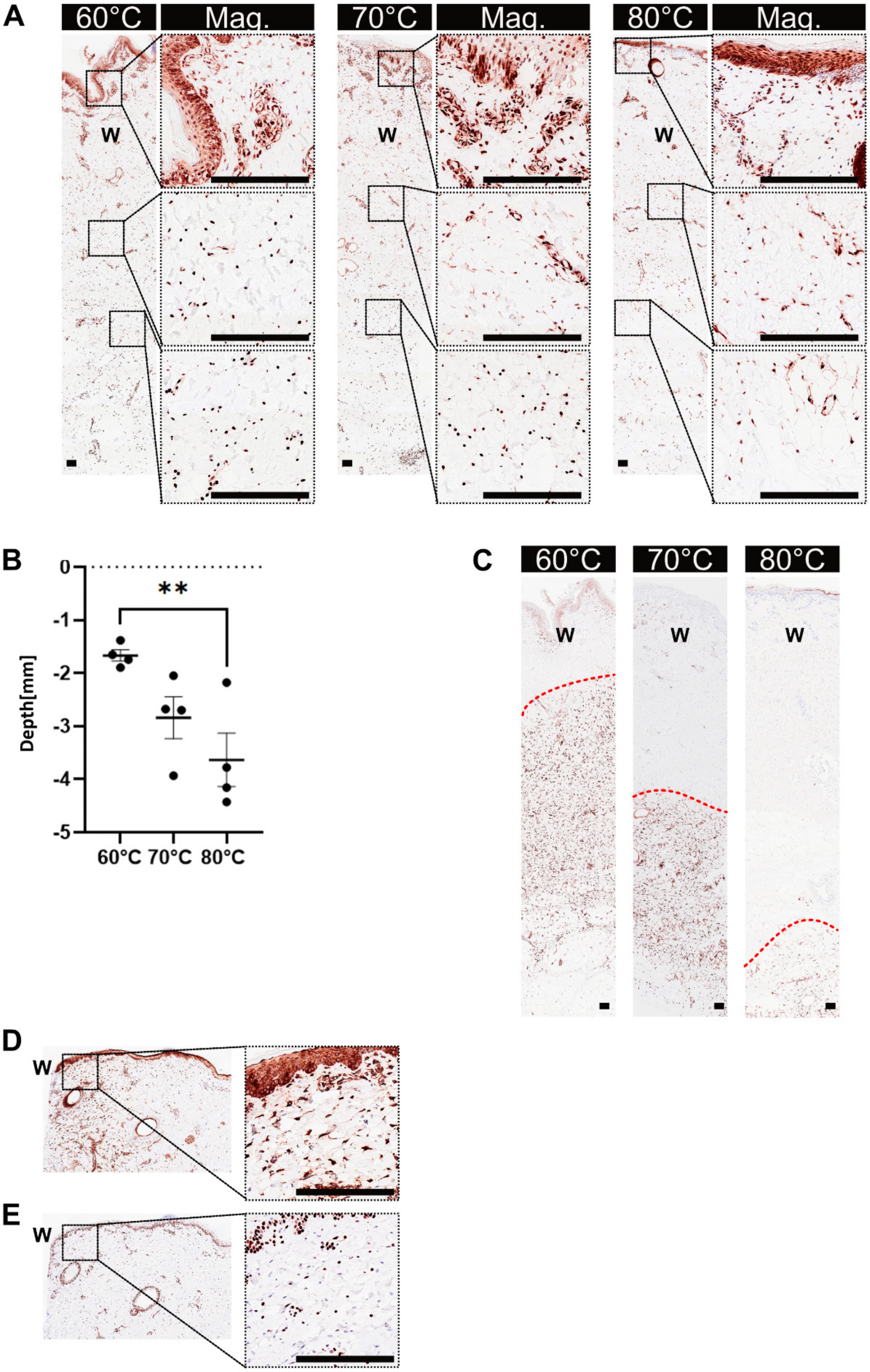
A
60°C
Mag.
70°C
Mag.
80°C
Mag.
W
W
W
B
0
-1
-2
-3
-4
-5
Depth[mm]
**
60°C 70°C 80°C
C
60°C
70°C
80°C
W
W
W
D
W
E
W

**Fig. S2. In burn injury the p-rpS6-zone stratifies cell death and survival responses.**

(**A**) Representative images of porcine skin biopsies collected 1.5 h after induction of 60, 70 and 80 ˚C burn injury and stained for HMGB1. Micrographs show magnified regions of epidermis (top panel) and two dermal regions (middle and bottom panels). (**B**) Quantification of the depth of termination of the HMGB1 zone in response to the 60, 70 and 80 ˚C burn injury at 1.5 h after the burn. (**C**) Images of porcine skin biopsies stained for p-rpS6 in 60, 70 and 80 ˚C burn injury (p-rpS6-zone marked with a red dashed line). (**D**) Representative image of p-rpS6 staining at the site of the excision wound. (**E**) Representative image of HMGB1 staining at the site of the excision wound. Data are from n = 4 pigs per group for (B). Mean ± SEM plotted. For (B) one-way ANOVA with post-hoc Dunnet's test was used. **p<0.01. Burn wounds are marked with a "W". All scale bars show 100 μm.

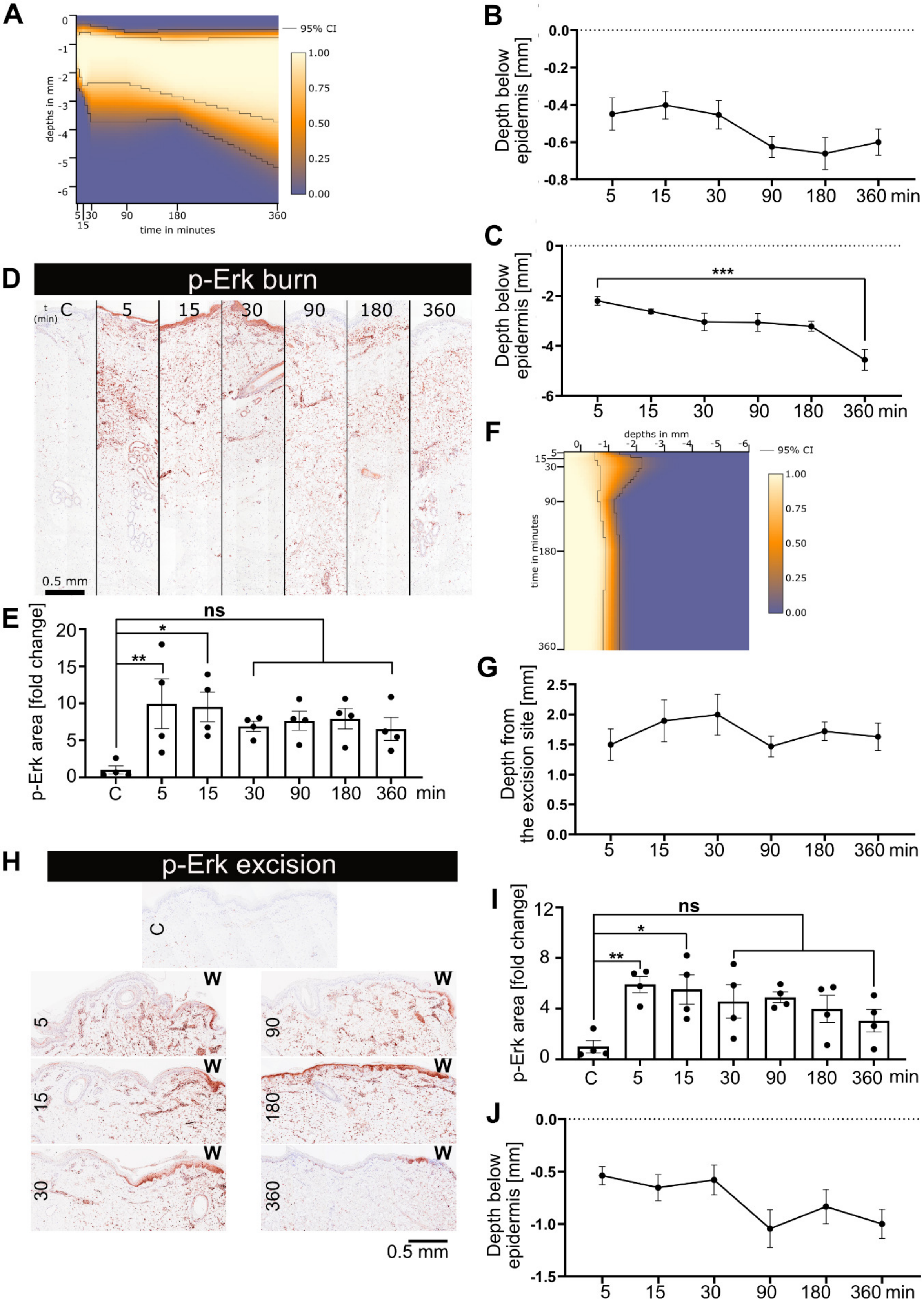
A
95% CI
depths in mm
time in minutes
0
-1
-2
-3
-4
-5
-6
1.00
0.75
0.50
0.25
0.00
5
15
30
90
180
360
B
Depth below epidermis [mm]
0.0
-0.2
-0.4
-0.6
-0.8
5
15
30
90
180
360 min
C
Depth below epidermis [mm]
***
0
-2
-4
-6
D
p-Erk burn
t (min)
C
5
15
30
90
180
360
0.5 mm
E
p-Erk area [fold change]
ns
*
**
20
15
10
5
0
C
5
15
30
90
180
360
min
F
depths in mm
time in minutes
95% CI
G
Depth from the excision site [mm]
2.5
2.0
1.5
1.0
0.5
0.0
H
p-Erk excision
C
W
5
15
30
90
180
360
0.5 mm
I
p-Erk area [fold change]
ns
*
**
12
8
4
0
J
Depth below epidermis [mm]
0.0
-0.5
-1.0
-1.5

**Fig. S3. The induction of the p-rpS6-zone is an early-onset and long-term response.**

(**A**) Probability chart showing depth of p-rpS6 layer in samples collected 5 min, 15 min, 30 min, 1.5 h, 3 h and 6 h after induction of 60 ˚C burn injury in pig skin. (**B**) Quantification of the initiation depth of the p-rpS6-zone in porcine skin samples collected 5 min, 15 min, 30 min, 1.5 h, 3 h and 6 h after induction of 60 ˚C burn injury in pig skin. (**C**) Quantification of the termination depth of the p-rpS6-zone in porcine skin samples collected 5 min, 15 min, 30 min, 1.5 h, 3 h and 6 h after induction of 60 ˚C burn injury in pig skin. (**D**) Representative images of immunohistochemical staining against p-Erk in control conditions and 5 min, 15 min, 30 min, 1.5 h, 3 h and 6 h after induction of 60 ˚C burn injury. (**E**) Quantification of the area of p-Erk in color-deconvolved images of control sample and at set timepoints after induction of 60 ˚C burn injury. (**F**) Probability chart showing depth of p-rpS6 layer is samples collected 5 min, 15 min, 30 min, 1.5 h, 3 h and 6 h after induction of excision injury in pig skin. (**G**) Quantification of the termination depth of the p-rpS6-zone in porcine skin samples collected 5 min, 15 min, 30 min, 1.5 h, 3 h and 6 h after induction of excision wound in pig skin. (**H**) Representative images of immunohistochemical staining against p-Erk in control conditions and 5 min, 15 min, 30 min, 1.5 h and 6 h after induction of excision wound. (**I**) Quantification of the p-Erk area in color-deconvolved images of control samples and at set timepoints after induction of excision wound. (**J**) Quantification of the termination depth of the HMGB1 zone in porcine skin samples collected 5 min, 15 min, 30 min, 1.5 h, 3 h and 6 h after induction of 60 ˚C burn injury in pig skin. Data are from n = 4 pigs per group for all the graphs. Mean ± SEM plotted. For all the graphs one-way ANOVA with post-hoc Dunnet's test was used. *$p<0.05$, **$p<0.01$, ***$p<0.001$ and "ns" is "non-significant". All scale bars show 500 µm.

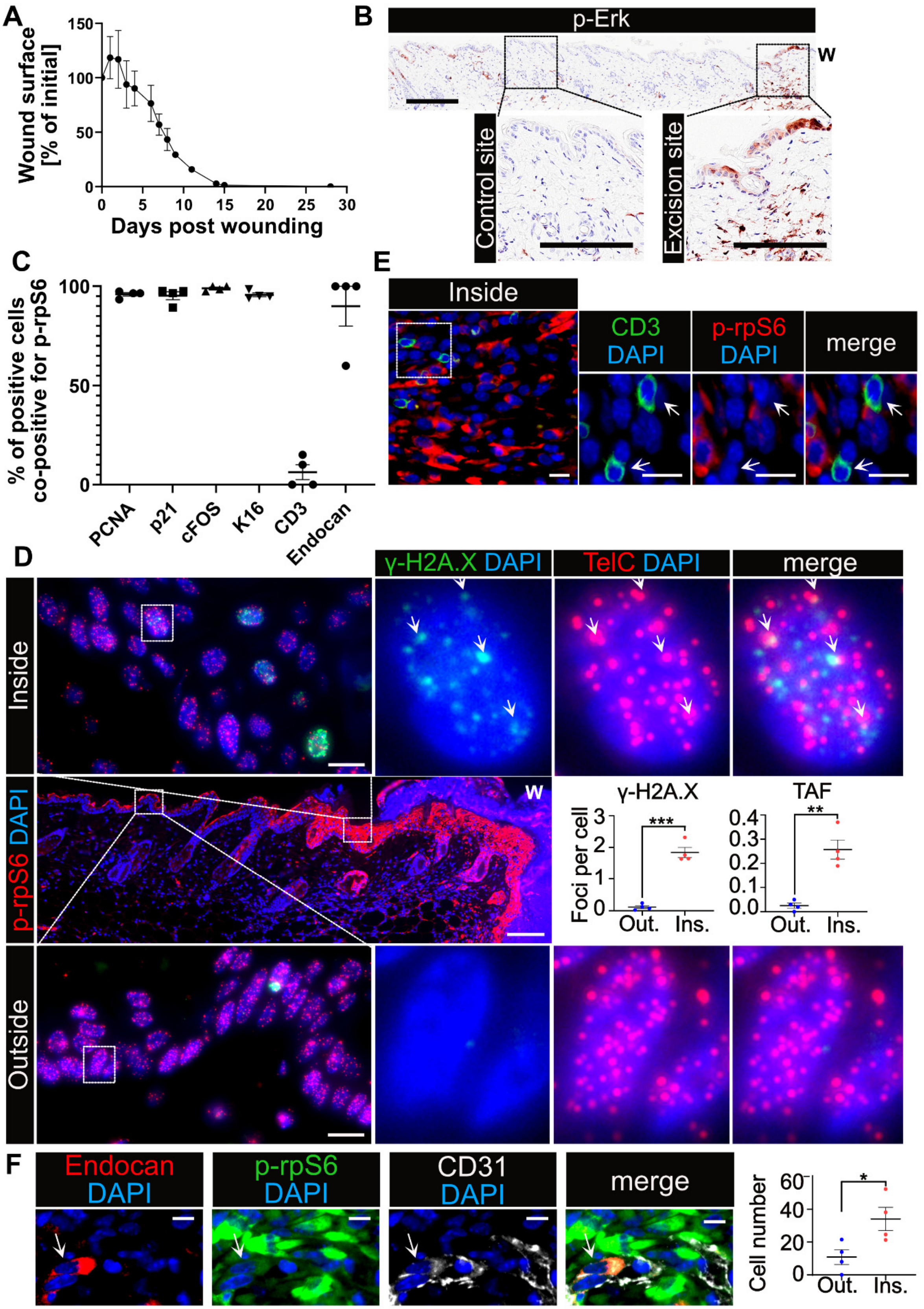

A
Wound surface [% of initial]
150
100
50
0
0 5 10 15 20 25 30
Days post wounding
B
p-Erk
W
Control site
Excision site
C
% of positive cells co-positive for p-rpS6
100
50
0
PCNA
p21
cFOS
K16
CD3
Endocan
E
Inside
CD3 DAPI
p-rpS6 DAPI
merge
D
γ-H2A.X DAPI
TelC DAPI
merge
Inside
p-rpS6 DAPI
W
γ-H2A.X
TAF
Foci per cell
***
**
3
2
1
0
0.4
0.3
0.2
0.1
0.0
Out.
Ins.
Outside
F
Endocan DAPI
p-rpS6 DAPI
CD31 DAPI
merge
Cell number
60
40
20
0
*
Out.
Ins.

**Fig. S4. The p-rpS6-zone is present throughout the healing process and encompasses pro-healing cellular processes.**

(**A**) Murine excision wounds were photographed at days 0, 1, 3, 4, 6, 7 8, 9, 11, 14, 15 and 28. Wound size was normalized to the measurement from day 0 and data is shown as percentage of initial size. (**B**) Representative images of immunohistochemical staining against p-Erk collected 30 min after excision injury of mouse skin. (**C**) Quantification of frequency of overlap of signals from the antibody against p-rpS6 and antibodies for PCNA, p21, cFos, K16, CD3 and Endocan. Data was analyzed in 5 images per mouse for PCNA, p21, cFos, and for K16, 4-10 images per mouse for CD3 where for all imaged were taken for regions inside the p-rpS6-zone and 14 images for Endocan. (**D**) Representative images of immunohistofluorescent stainings of the p-rpS6-zone and telomere DNA damage in successive sections of murine skin collected 3 days after excision injury. The middle image shows the whole wound area, stained forp-rpS6 (red) and co-stained with DAPI (blue). Above and below there are close-ups of areas inside and outside the p-rpS6-zone, stained for TelC (red), γ-H2A.X (green) and co-stained with DAPI (blue). Micrographs show individual cells. White arrows mark telomere associated foci (TAF). Graphs show quantification of average number of γ-H2A.X foci and TAF per cell present outside (Out.) or inside (Ins.) the p-rpS6-zone. 6 images comprising 200 cells per mouse were quantified. (**E**) Representative images of immunohistofluorescent staining against p-rpS6 (red), CD3 (green) and co-stained with DAPI (blue) in a sample collected 3 days after excision injury of mice. Micrographs show regions inside the p-rpS6-zone. White arrows mark CD3-positive cells. (**F**) Representative images of immunohistofluorescent staining against p-rpS6 (green), endocan (green) and co-stained with DAPI (blue) in a sample collected 12 days after excision injury of mice. White arrows mark an endocan-positive cell. Graph shows quantification of average number of cells positive for endocan and present outside (Out.) or inside (Ins.) the p-rpS6-zone. 14 images per mouse were quantified. Data are from n = 4 mice per group for all the graphs. Mean ± SEM plotted for all the graphs. Scale bars for (B), the macrographs of (D) and (E) show 100 μm, for the micrographs in (D), (E) and (F) scale bars are 10 μm. *$p<0.05$, **$p<0.01$, ***$p<0.001$.

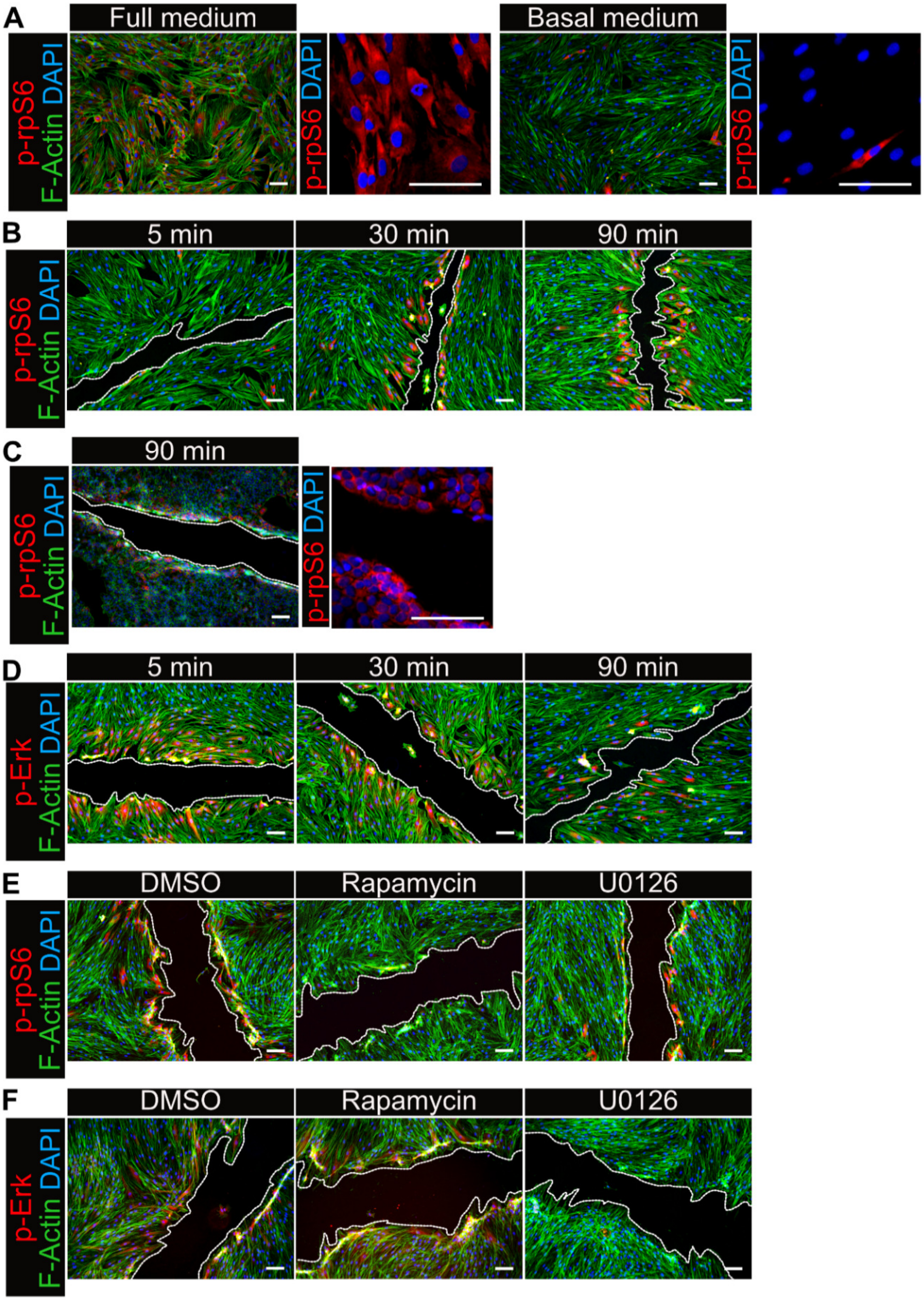
A
Full medium
Basal medium
p-rpS6 F-Actin DAPI
p-rpS6 DAPI
B
5 min
30 min
90 min
p-rpS6 F-Actin DAPI
C
90 min
p-rpS6 F-Actin DAPI
p-rpS6 DAPI
D
5 min
30 min
90 min
p-Erk F-Actin DAPI
E
DMSO
Rapamycin
U0126
p-rpS6 F-Actin DAPI
F
DMSO
Rapamycin
U0126
p-Erk F-Actin DAPI

**Fig. S5. Formation of the p-rpS6-zone is induced by DAMPs and dependent on mTOR.**

(**A**) Representative images of immunofluorescent staining against p-rpS6 (red) and co-stained with phalloidin-488 (marking F-Actin; green) and DAPI (blue) in human dermal fibroblasts (HDFs) kept in full media (containing foetal bovine serum; FBS) or basal media (without FBS). (**B**) Representative images of immunofluorescent staining against p-rpS6 (red) and co-stained with phalloidin-488 (green) and DAPI (blue) in HDFs fixed 5, 30 or 90 min after induction of the scratch assay. (**C**) Representative images of immunofluorescent staining against p-rpS6 (red) and co-stained with phalloidin-488 (green) and DAPI (blue) in HaCat cells fixed 5, 30 or 90 min after induction of the scratch assay. (**D**) Representative images of immunofluorescent staining against p-Erk (red) and co-stained with phalloidin-488 (green) and DAPI (blue) in HDFs fixed 5, 30 or 90 min after induction of the scratch assay. (**E**) Representative images of immunofluorescent staining against p-rpS6 (red) and co-stained with phalloidin-488 (green) and DAPI (blue) in HDFs treated with DMSO, Rapamycin or U0126 and fixed 30 min after induction of the scratch assay. (**F**) Representative images of immunofluorescent staining against p-Erk (red) and co-stained with phalloidin-488 (green) and DAPI (blue) in HDFs treated with DMSO, Rapamycin or U0126 and fixed 30 min after induction of the scratch assay.

All scale bars show 100 µm.

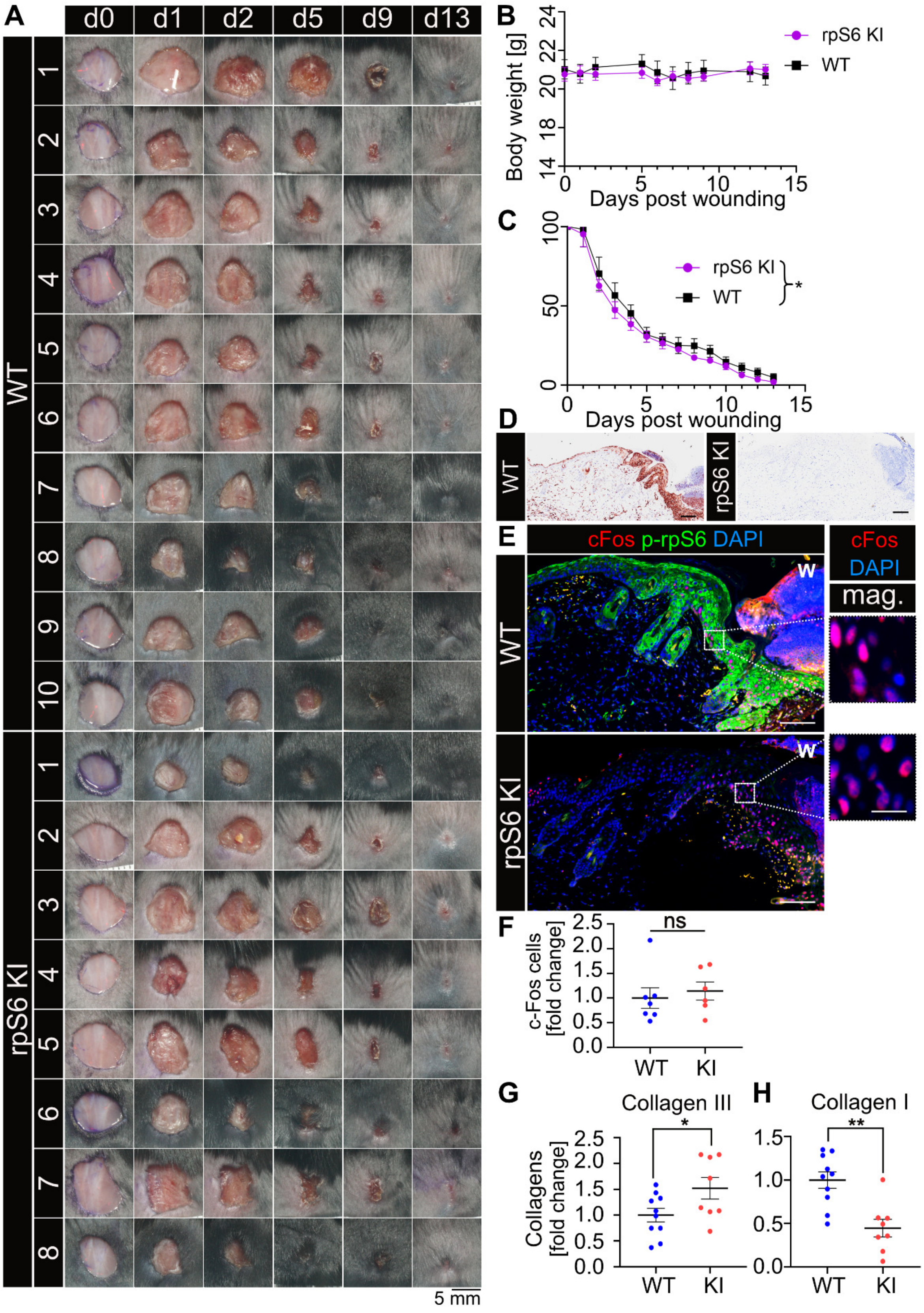
A
d0
d1
d2
d5
d9
d13
WT
rpS6 KI
1
2
3
4
5
6
7
8
9
10
5 mm
B
Body weight [g]
Days post wounding
rpS6 KI
WT
C
Days post wounding
rpS6 KI
WT
*
D
WT
rpS6 KI
E
cFos p-rpS6 DAPI
cFos
DAPI
mag.
W
F
c-Fos cells [fold change]
ns
WT
KI
G
Collagen III
Collagens [fold change]
*
H
Collagen I
**

**Fig. S6. P-rpS6 deficiency results in faster initial wound closure but disrupted healing.**

(**A**) Photographs of WT and rpS6 KI wounds throughout the process of healing. (**B**) Average body mass of WT and rpS6 p-/- KI (rpS6 KI) mice throughout the process of wound healing from day 0 to day 13 post wounding. (**C**) rpS6 KI and WT littermate mouse excision wounds were photographed at days 0-13 after wounding. Wound size was normalized to the measurement from day 0 and data is shown as percentage of initial size. (**D**) Representative p-rpS6 staining in skin collected from WT and rpS6 KI mice at 3 d post injury. (**E**) Murine skin at 3 d post injury, with micrographs (without p-rpS6 staining) and quantifications of the region immediately adjacent to the wound in rpS6 KI and WT male mice. cFos (red), p-rpS6 (green) and DAPI (blue). (**F**) Quantification of average number of cFos-positive keratinocytes. (**G**) Quantification of collagen III level in skin of rpS6 KI and WT mice. (**H**) Quantification of collagen I level in skin of rpS6 KI and WT mice. Data are from n = 8 female WT and n = 10 female rpS6 KI mice for (A) and (B), from n = 9 male WT and n = 8 male rpS6 KI littermates for (C), and from n = 6 male WT and n = 7 male rpS6 KI mice for (F). Mean ± SEM plotted for all the graphs. For the graphs (F), (G) and (H) unpaired t test was used. The scale bar for the skin photographs (A) is 5 mm, for the images in (D) and (E) it is 100 µm and for the micrographs 10 µm. *p<0.05, **p<0.01 and “ns” is “non-significant”.

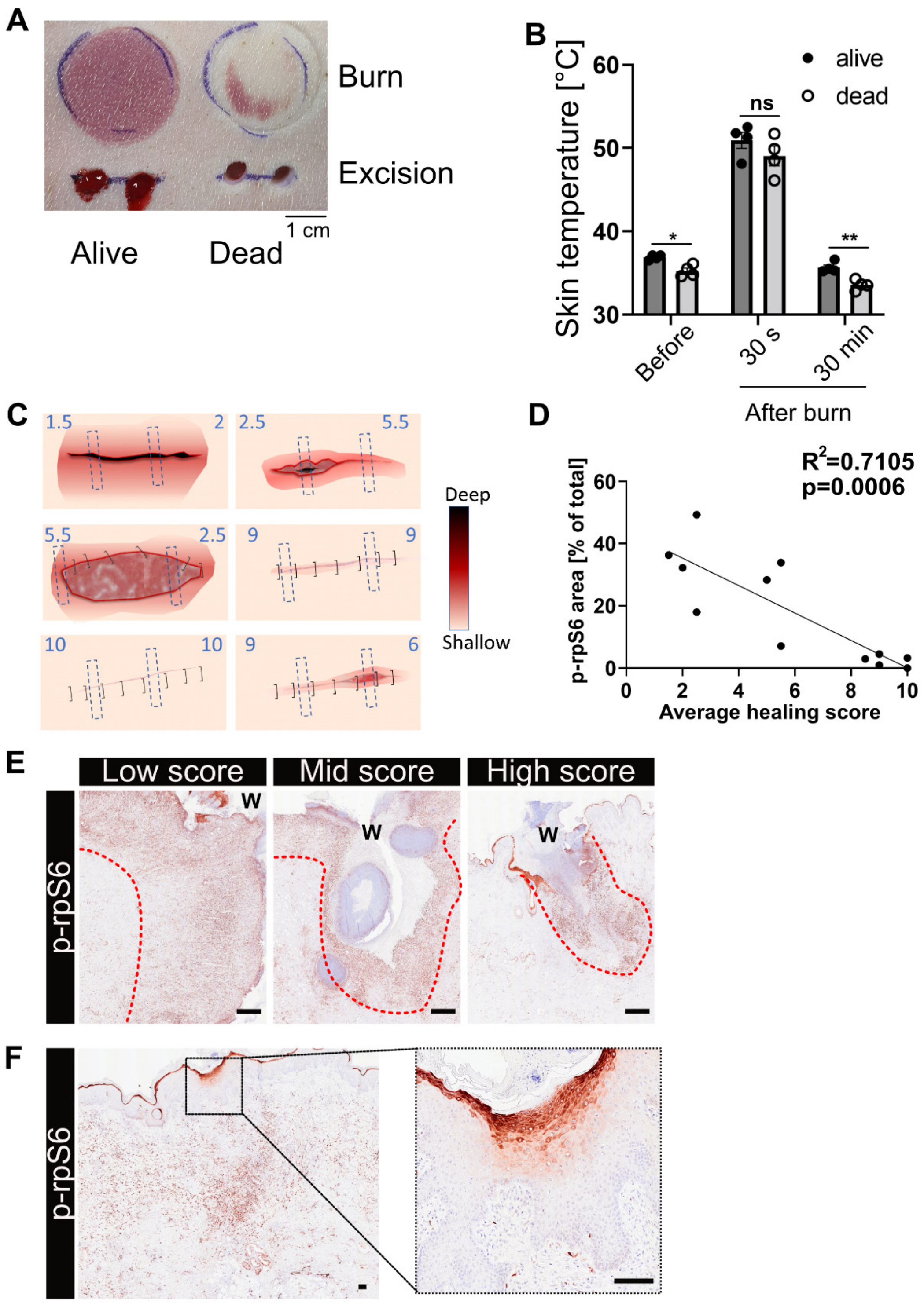
A
Burn
Excision
1 cm
Alive
Dead
B
Skin temperature [°C]
alive
dead
ns
*
**
Before
30 s
30 min
After burn
C
Deep
Shallow
D
$R^2$=0.7105
p=0.0006
p-rpS6 area [% of total]
Average healing score
E
Low score
Mid score
High score
p-rpS6
W
F
p-rpS6

**Fig. S7. The p-rpS6-zone reports on oxygen availability, vascularization, and progression of healing in pre-clinical studies.**

(**A**) Photography of pig skin showing erythema caused by a burn injury performed when the animal was alive compared to the lack of erythema when burned at 3 min post-death. (**B**) Measurements of skin temperature taken prior to burn injury, 30 s after the injury or 30 min later. (**C**) Schematics that are graphical representations of the porcine incision injuries 7 days post wounding. The blue lines represent the region from which the biopsy samples were collected (and thus also the histological orientation of the subsequent sections). The number next to the line is an average of the wound assessment score as assigned independently by two trained veterinarians (where 1 represents an unhealed wound and 10 represents a fully healed wound).

(**D**) Graph showing correlation between the wound assessment score and the area of p-rpS6-positive signal when normalized to the total area of each section. (**E**) Porcine skin collected 7 days after wounding with low (left panel), medium (middle panel) and high (right panel) wound assessment scores, stained for p-rpS6. Only one side of the wound with the lowest score is shown. (**F**) A representative image of an incision wound with one of the highest wound scores (most healed) that still shows remnants of p-rpS6 in epidermis and dermis even after the apparent macroscopical completion of the healing process.

Data are from n = 4 pigs for (B). Two samples were collected from each incision wound from 6 pigs at 7 days post injury resulting in n = 12 biological replicates for the graph (D). Mean ± SEM plotted for (B). For the graph (B) two-way ANOVA with post-hoc Sidak's test was used. For the graph (D) Pearson's correlation test was used. The scale bar for the skin photographs (B) is 1 cm and for images (E) and (F) it is 100 μm. *$p<0.05$, **$p<0.01$ and "ns" is "non-significant".

**Movie S1.**

The sample of porcine needle prick sample was sectioned horizontally starting from the epidermis into 40 slices of 4 μm thickness with 20 μm gaps between each slice. All 40 slices were stained for p-rpS6, acquired on a scanning microscope and then used to create a 3D projection representing regions of tissue positive for p-rpS6 staining. The 3D projection shows the needle (green) and tissue positive for p-rpS6 staining (red) above epidermis (cyan).
